# An agent-based model of wound healing in fibroblast scratch assays

**DOI:** 10.64898/2026.10.15.759485

**Authors:** Rowan Sumanaweera, Cooper Stansbury, Walter Meixner, Jillian Cwycyshyn, Lindsey Muir, Indika Rajapakse

## Abstract

Wound healing assays are widely used to study fibroblast migration and proliferation in the lab. Although traditional scratch assays provide valuable insight, they can be time-intensive, inflexible, and difficult to reproduce. Computational models provide an efficient and reproducible way to study fibroblast behavior in silico. We developed an agent-based simulation of fibroblast scratch assays that models migration, proliferation, contact inhibition, cell-cell mechanical interactions, and the extracellular matrix (ECM) through a particle representation of fibroblasts and the ECM. To validate our model, we performed two tests: a leave-one-out test, where model parameters were calibrated to 6 imaging time series and tested on the remaining one, and a prediction test, where model parameters were calibrated to the first 24 hours of data and the remaining 66 hours of data were predicted. In our leave-one-out test, our model achieved a mean normalized root mean squared error (mean NRMSE) of 3.34%, and a Pearson’s R of *>* 0.98 across all tested wound geometries time series. In the prediction test, the model predicted the remaining 66 hours of wound closure with an NRMSE of 2.08%. These results demonstrate that experimentally observed wound healing behavior of fibroblasts across different geometries can be modeled by local, agent-based rules. Our model provides an efficient platform for the experimentation and in silico study of fibroblast behavior.

## 1 Introduction

Wound healing restores the structural and functional integrity of organs and tissues after injury. Problems during wound healing can cause persistent inflammation, fatal bacterial infections, and long-lasting scars. Fibroblasts, the most common stromal cell type, play a central role in tissue regeneration and the rebuilding of the extracellular matrix (ECM) during wound healing. To better understand fibroblast behavior, 2D wound healing assays such as scratch assays or barrier-based assays are commonly used [17, 19]. These traditional methods are simple and effective, but can be time-intensive, inflexible, and difficult to reproduce. Computational models of scratch assays can allow many flexible, reproducible experiments to be run in parallel with precise control over model parameters.

To facilitate the study of wound closure, we developed an agent-based computational model of an invitro scratch assay with cell-cell repulsion forces, interactions with the ECM, and control over wound geometry. The model provides an efficient framework for studying how fibroblast proliferation and migration affect wound healing while enabling easy comparison with experimental data. To evaluate our model, we compared simulated wound closure dynamics to experimentally observed data across multiple wound geometries, and quantified goodness of fit using several metrics. This allowed us to draw conclusions about the model’s general ability to replicate experimental data, rather than a single wound shape.

There have been several approaches to modeling wound healing: continuum-based models, hybrid models, and agent-based models. Tranquillo and Murray (1992) [20] developed one of the first continuum-based models of dermal wound contraction, representing fibroblasts, the ECM, and inflammatory mediators as fields governed by partial differential equations that model tissue dynamics. Olsen et al. (1995) [16] continued this approach, including mechanochemical interactions between fibroblasts and tissue.

Dallon et al. (1999) [7] later developed one of the first hybrid models, tracking discrete fibroblast positions and treating the ECM as a continuous field. Unlike our model, their work primarily focused on dermal healing by first intention (in which the edges of a wound are held closely together) and the evolution of ECM fiber organization and alignment.

As computing power grew in the early 2000s, researchers adopted fully agent-based approaches to better model the intricacies of wound healing. One of the earliest fully agent-based models—Walker et al. (2004) [21]—demonstrated that tissue-level dynamics could emerge from local cell-level interactions. Because agent-based models represent fibroblasts explicitly, they are well-suited for studying how migration, proliferation, and contact inhibition directly influence wound closure.

In contrast to many existing models, our simulation represents both fibroblasts and the ECM as discrete particles, and its large-scale dynamics emerge from various interactions between cells and between cells and ECM, such as mechanical repulsion and ECM avoidance. Our explicit modeling approach, in combination with our model’s focus on fibroblast scratch assays, allows us to investigate wound healing dynamics at both the population level and the agent level.

## 2 Methods

### 2.1 Model Overview

A complete, detailed model description, following the Overview, Design concepts, Details (ODD) protocol (Grimm et al. 2006, 2010, 2020) [10, 11, 12] is included in Appendix B. The following is a summary of that document.

This model proposes the wound healing behavior of fibroblasts can emerge from simple local rules such as migration, proliferation, contact inhibition, mechanical crowding, and interactions with the extracellular matrix (ECM). The purpose of the model is to predict the dynamics of fibroblast proliferation and migration in a monolayer scratch assay across different wound configurations to reduce the cost, time, and reproducibility problems faced by conventional scratch assays. To validate the model, we use the following patterns: a) wound area reduction over time—the model should reproduce the different wound area-time curves observed in the test data across different wound geometries, and b) healing times—the model aims to reproduce the experimentally observed time to 98% closure of our test data.

The model includes the following entities: a) fibroblasts, the primary agent, and b) ECM particles, non-agentic representations of the extracellular matrix. For each simulation step, each fibroblast agent executes a series of processes which govern its behavior: the physics submodel, the locomotion submodel, the cell cycle submodel, and the ECM deposition submodel. Each fibroblast agent executes these processes in sequence, and the fibroblast agents are updated in an arbitrary order. An overview of the processes is shown in Figure 1.

**Figure 1:**
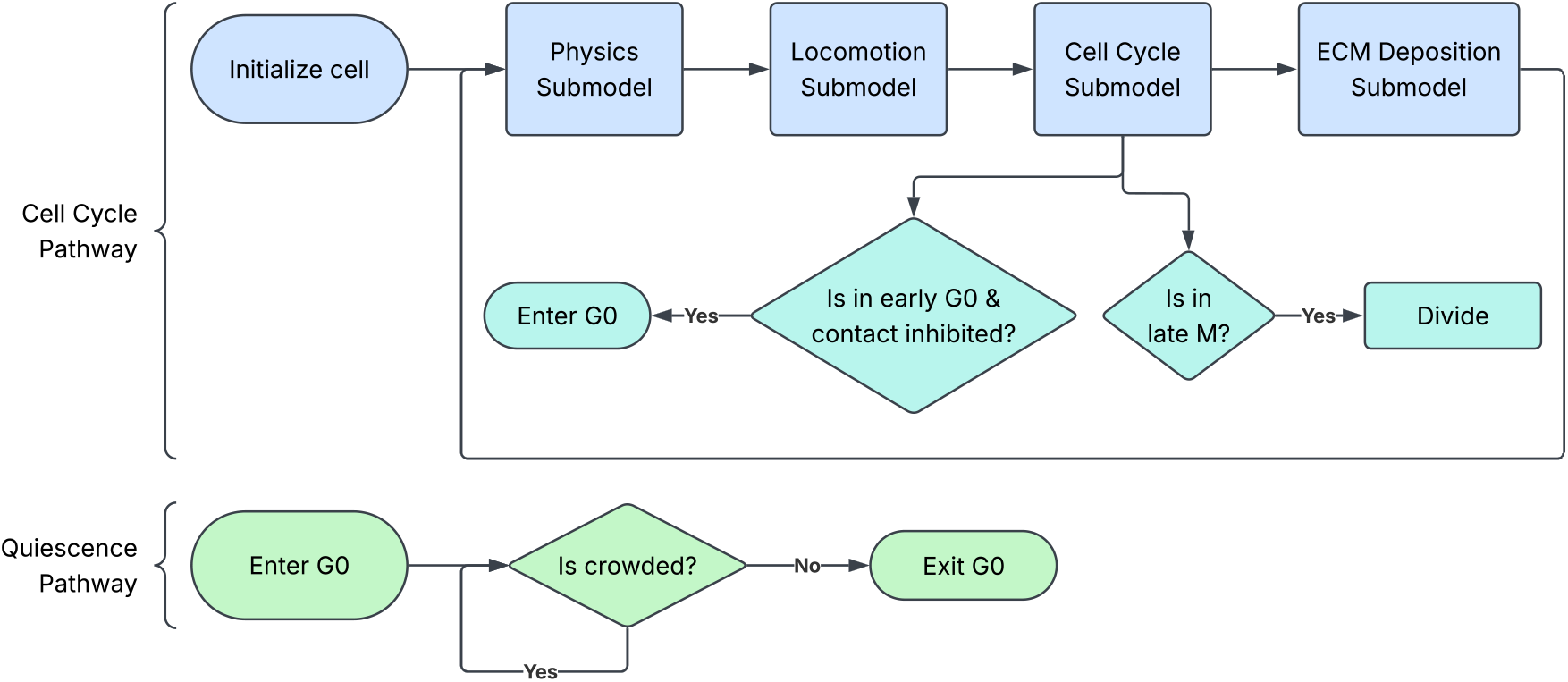
An overview of the submodels executed for each fibroblast every simulation step, for both the (upper) cell cycle pathway, and the (lower) quiescence pathway.

The processes are described as follows:

1. Physics. The agent’s position is updated based on its previous position, and physical constraints such as collision handling and boundary conditions are applied. Forces applied by this submodel help create a uniform monolayer, and in overcrowded areas of a monolayer, those forces play a key part in pushing fibroblasts into the wound.
2. Locomotion. The agent’s turn-state variable is updated stochastically, and its position is updated based on local ECM density and its current speed and turn-state variables. Once a wound is formed, the fibroblast agent migrates inwards in a Correlated Random Walk (CRW) [3, 5], away from high local concentrations of ECM.
3. Cell cycle. The agent’s cycle phase is advanced according to a configured timer. When a cell exits the M phase, a new fibroblast agent is created at the parent cell’s position. Cells can also enter contact inhibition at specific checkpoints if the local density of fibroblasts exceeds a certain threshold.
4. ECM deposition. An ECM particle is deposited at the agent’s position according to a configured, varying timer. In high concentrations of ECM, the deposition rate slows to limit runaway growth, and in low concentrations, it speeds up.

The spatial resolution of the simulation was set to 8.2mm x 8.2mm for the purposes of this publication, but is variable. The temporal resolution is also adjustable but is fixed to 2.5 hours per step for this publication. All time-related parameters (e.g., cycle duration) are expressed in simulation steps.

The most important design concepts of the model are discrete fibroblast representation, response to ECM, and emergence-driven wound closure. The model follows an agent-based modeling approach that explicitly represents each fibroblast as a circle, in contrast to a continuum-based model [1] or Cellular Potts model [9]. Cell locomotion is heavily influenced by ECM particles, and during the cell cycle submodel, the cells deposit ECM particles, creating a feedback loop which encourages cells to migrate into areas with low fibroblast densities. One of the most fundamental properties of the simulator is the implicit definition of wound closure. No simulation rules explicitly define a wound, wound boundary, or wound closure objective. Rather, fibroblasts exhibit wound healing behavior as an emergent byproduct of the built-in proliferation, migration and reaction to ECM.

The model incorporates fibroblast migration and proliferation, contact inhibition, mechanical forces, and response to the ECM because these processes directly affect the spatial redistribution of fibroblasts and temporal healing rates during closure. Migration allows cells to move into the wound, while proliferation allows the cells to fill it. Contact inhibition and mechanical forces prevent cells from overcrowding and regulate their population, and interactions with the ECM allows cells to influence each other’s migration. We selected these processes as a set of minimal rules that could be capable of producing the large-scale dynamics observed in wound healing. That goal led us to represent both cells and ECM as points as a simplification deemed acceptable to model wound closure dynamics. The simulation does not attempt to model every cell-level process of wound healing; instead, it aims to replicate the macroscopic spatial and temporal patterns of wound closure.

### 2.2 Model Validation

#### 2.2.1 Data

We validated our method by comparing our simulation to several imaging data time series. These time series were captured with a Zeiss CellDiscoverer7 microscope fitted with an Axiocam 506 camera by Cwycyshyn et al. [6], using human BJ fibroblasts. The dataset consisted of 7 videos with each having different wound geometries (2 circles, 1 line, 1 square, and 3 triangles). In each trial, a uniform cell layer was grown in vitro, and a specific wound shape was applied by the BioAssemblyBot robotic platform [6]. Each time series is approximately 90 hours in length, of 271 total frames, with each frame covering an 8.2mm x 8.2mm area.

For each trial, we extracted the measured wound area vs. time curves from the data, *A*(*t*), and the predicted wound area from our simulator, *a*(*t*), and compared the two using several figures-of-merit to evaluate our simulation’s accuracy under different initial areas and geometries.

#### 2.2.2 Wound Area Estimation

We extracted the wound area from the experimental data and the simulated data using two distinct methods.

##### Experimental Data

The microscope, using oblique illumination, assembles several tiles to produce each video frame as shown in Figure 2(a). There were image artifacts at the tile boundaries. We high pass filtered each frame using a Gaussian filter kernel to produce the intermediate frames shown in Figure 2(b). The texture in the intermediate frames are first quantified using the local standard deviation to produce a series texture frames.

**Figure 2:**
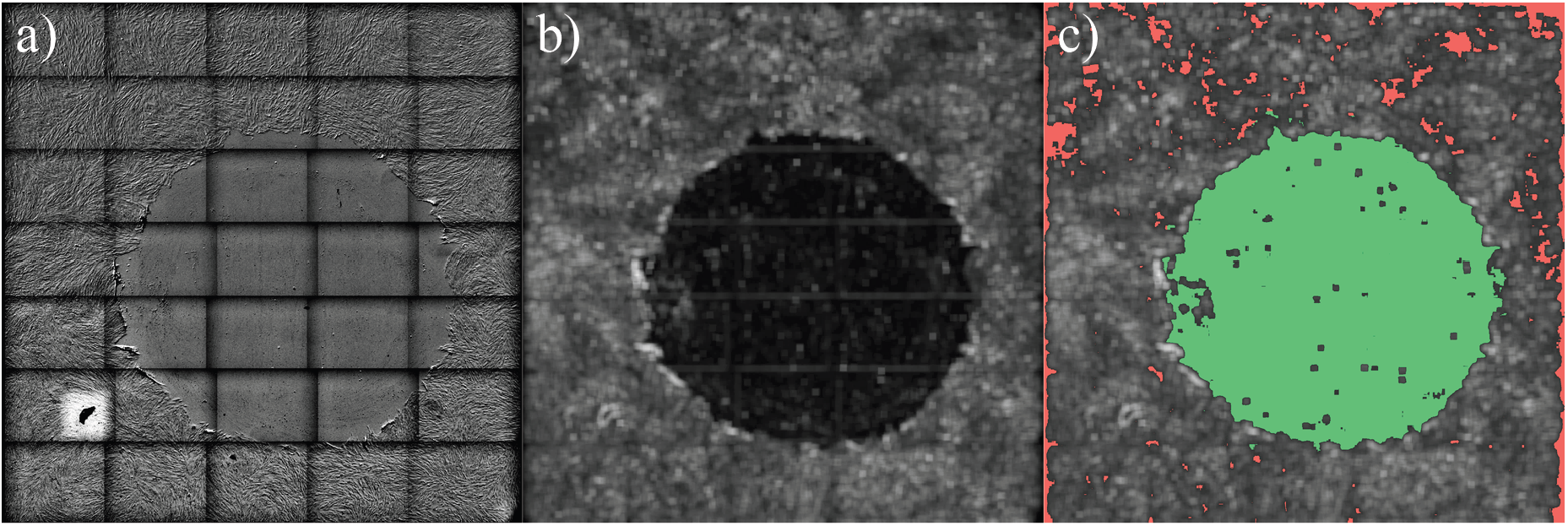
Representative snapshots from the Circle 1 dataset at selected points of our wound area estimation process: a) before imaging processing, b) after high pass filtering, and c) after thresholding and edge detection (red indicates spurious pixels, while green indicates non-spurious).

Next, pixels with texture values greater than a threshold, T, are tagged as “wound pixels”, and all others are tagged as normal tissue. The value of T is determined by examining the bimodal histogram of all pixels in a given video. T corresponded to the approximate minimum value between the two peaks of the bimodal histogram. There were spurious pixels outside the wound area as well (Figure 2(c)). We used edge detection followed by dilation to produce a mask to exclude such spurious pixels outside the wound area.

##### Simulated Data

To replicate the initial wound shape of the experimental data, we chose to create our simulated wound by removing cells within the area of an idealistic geometric shape. We fit the area of the shape (circle, square, line, or triangle) to the initial area of the experimental data.

To estimate the wound area throughout a simulation run from the simulated data, the simulation space is divided into a grid where each grid stores the fibroblast occupancy count *C*(*x, y*). Then, a binary mask *M* (*x, y*) is created by

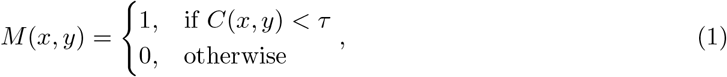

where *τ* is a fixed threshold on fibroblast occupancy.

Then, the simulated area *A*(*t*) is calculated by summing over all grid cells classified as wound by the mask as defined by

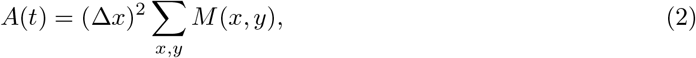

where Δ*x* is the length of one grid cell.

#### 2.2.3 Tests

We tested the ability of our simulator to predict wound healing using two separate tests:

1. Leave-one-out Test: The 7 videos were divided into 6 training videos and 1 testing video. For each trial, We used the training dataset to estimate the 7 simulation parameters by fitting the simulation model to the data by minimizing the objective function (see Section 2.2.4). The goodness-of-fit for each trial was quantified using 3 metrics:
  a. NRMSE: The RMS error normalized with the starting wound area expressed as a percentage.
  b. *R*: Pearson’s correlation coefficient between the prediction, *a*(*t*) and the measurement, *A*(*t*).
  c. Δ*T*_*c*_: The error of the time for the wound to close 98% of the initial area, between the prediction and the measurement. We also report a normalized Δ*T*_*c*_, defined by 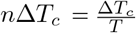, where *T* is the total simulation time duration.
2. Prediction Test: Here we divided each waveform into 2 sections: (a) first 24 hours, (b) remaining 66 hours. We then tuned the selected simulation parameters to minimize Φ using the first 24 hours, and predicted the wound healing process for the remaining 66 hours. We quantified the goodness of fit for each waveform with the same metrics as the leave-one-out trial (NRMSE, R, Δ*T*_*c*_, *n*Δ*T*_*c*_). These metrics were computed over only the final 66 hours, based on the predictions generated from the first 24 hours.

We selected these metrics because each metric alone cannot capture all aspects of the agreement between the waveforms. For example, RMSE and R are simply measures of agreement, while Δ*T*_*c*_ indicates whether the simulation predicted closure too late or too early. Together, these metrics allow us to understand not only the agreement between the waveforms, but also the lag time of closure.

#### 2.2.4 Optimization

To minimize the objective function for our tests, we selected the following 7 parameters to optimize: *cell-repulsion, max-speed, turn-speed, turn-chance, inhibition-radius, ecm-avoidance*, and *ecm-speed-scalar*. Let *θ* = the vector of all optimization parameters. We bounded each parameter as shown in Table 1.

**Table 1:** Parameter bounds used in optimization and sensitivity analysis.

| Parameter | Lower Bound | Upper Bound |
| --- | --- | --- |
| cell-repulsion | 0.1 | 0.5 |
| max-speed | 0.5 | 5.0 |
| turn-speed | 0.1 | 1.0 |
| turn-chance | 0.1 | 0.9 |
| inhibition-radius | 1.8 | 1.9 |
| ecm-avoidance | 0.1 | 10.0 |
| ecm-speed-scalar | 0.1 | 30.0 |

For this procedure, we used Gaussian process-based Bayesian optimization implemented by the gp minimize function in the Python library Scikit-optimize. We selected this method because each simulation run is computationally expensive, and Bayesian optimization more efficiently explores parameter spaces compared to alternatives, such as an exhaustive search of the parameter space. For each optimization run, we set the number of random initial points to 15, and the number of total simulation runs to 100. The simulation was run once per objective function evaluation, which generated *a*(*t*).

For all of our optimization tests, we selected the Root Mean Squared Error (RMSE) as the metric for goodness of fit, defined by:

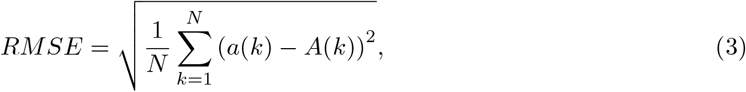

where *N* is the number of sampled time points.

For our leave-one-out test, we picked one dataset to leave out, and trained the model on the other six. We tested the optimized model on the omitted dataset, reported the results, and repeated this process for all seven datasets. For the leave-one-out optimization process, we defined the objective function Φ(*θ*) as the sum of the RMSE between the predicted wound area function *a*(*t*) and the measured function *A*(*t*) of each training dataset, as defined by:

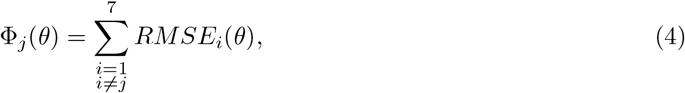

where the omitted dataset is denoted by *j*.

For our prediction test, we trained the model on only the first 24 hours of the run, and then tested the optimized model on the last 66 hours, and reported the results. For the prediction optimization process, we defined the objective function Φ(*θ*) as the RMSE restricted to the domain *t* ∈ [0, 24].

#### 2.2.5 Sensitivity Analysis

In addition to the above tests, we performed a global sensitivity analysis using the Morris method [13], which uses the one-factor-at-a-time approach. This method isolates a single parameter per run, and varies it to determine parameter significance. In our implementation, we used the SALib Python library. We report the *µ*^∗^ metric proposed by Campolongo et al. [4] to report the significance of each parameter.

We used the same set of parameters as in the optimization procedure, with the same bounds for consistency, shown in Table 1. The objective function was also defined as the RMSE between the real and simulated curves for the procedure. We performed the sensitivity analysis using 12 Morris trajectories and 6 discretization levels per parameter. The parameter space was normalized to [0, 1]^*n*^, and each trajectory consisted of *n* + 1 model evaluations, where *n* is the number of parameters.

#### 2.2.6 Spatial Analysis

In order to extend our conclusions beyond the temporal, but not spatial, system-level data of the Leave-one-out and Prediction tests, we mapped the spatial migration and proliferation dynamics of the optimized models from both tests.

We divided the simulation space and real video data into a 16x16 grid, and measured a wound area time series individually for each grid cell. Then, we generated two heatmaps, one using RMSE to compare the two time series and the other using Δ*T*_*c*_.

## 3 Results

Figure 3 compares our visual simulation output to both the raw and processed training data at selected time points.

**Figure 3:**
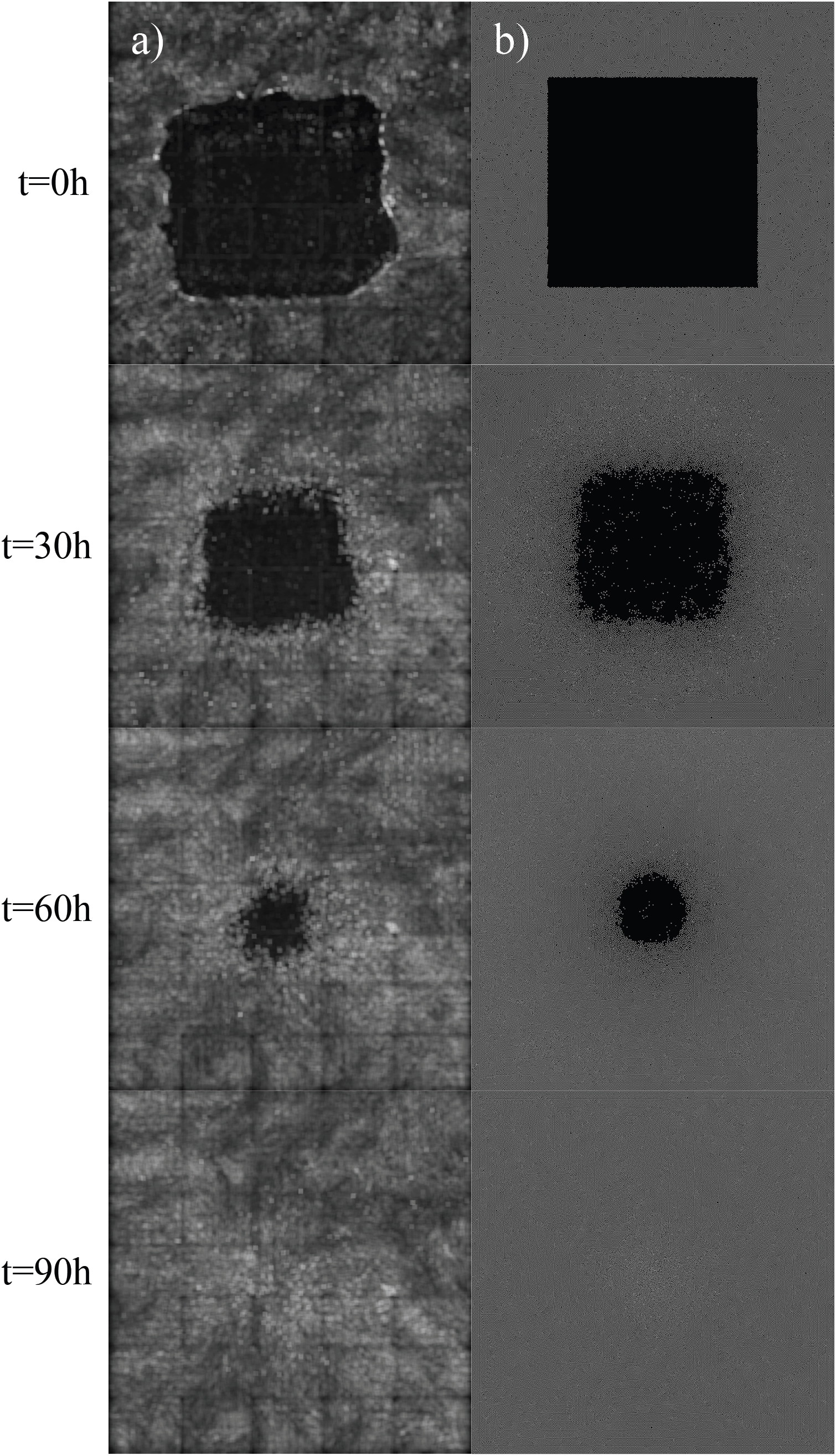
Representative snapshots from the Square dataset at selected time points of a) our processed image data following the steps outlined in Section 2.2.2, and b) visual output from the simulation.

### 3.1 Leave-one-out Test

Figure 4 shows *a*(*t*) overlaid with *A*(*t*) for each of the Circle 1, Square, Line, and Triangle 1 datasets after tuning the simulator using the remaining 6 test videos. Table 2 lists NRMSE, *R*, Δ*T*_*c*_, and *n*Δ*T*_*c*_ for each of the test videos. The mean NRMSE was 3.34%, the Pearson’s *R* was greater than 0.98 for all cases, and *n*Δ*T*_*c*_ was under 10% for 6 of the 7 videos.

**Table 2:** Mean leave-one-out model performance across datasets.

| Dataset | NRMSE | R | $\Delta T_c$ (h) | $n\Delta T_c$ |
| --- | --- | --- | --- | --- |
| Circle 1 | 2.25% | 0.9992 | 5.41 | 6.01% |
| Circle 2 | 3.95% | 0.9981 | 6.38 | 7.09% |
| Square | 3.07% | 0.9986 | 2.04 | 2.26% |
| Line | 2.55% | 0.9959 | -0.06 | 0.06% |
| Triangle 1 | 2.60% | 0.9992 | -7.31 | 8.12% |
| Triangle 2 | 4.33% | 0.9980 | -7.90 | 8.78% |
| Triangle 3 | 4.65% | 0.9974 | -16.28 | 18.09% |
| $\mu$ | 3.34% | 0.9981 | -2.53 | 2.81% |
| $\sigma$ | 0.96% | 0.0011 | 8.27 | 9.19% |

**Figure 4:**
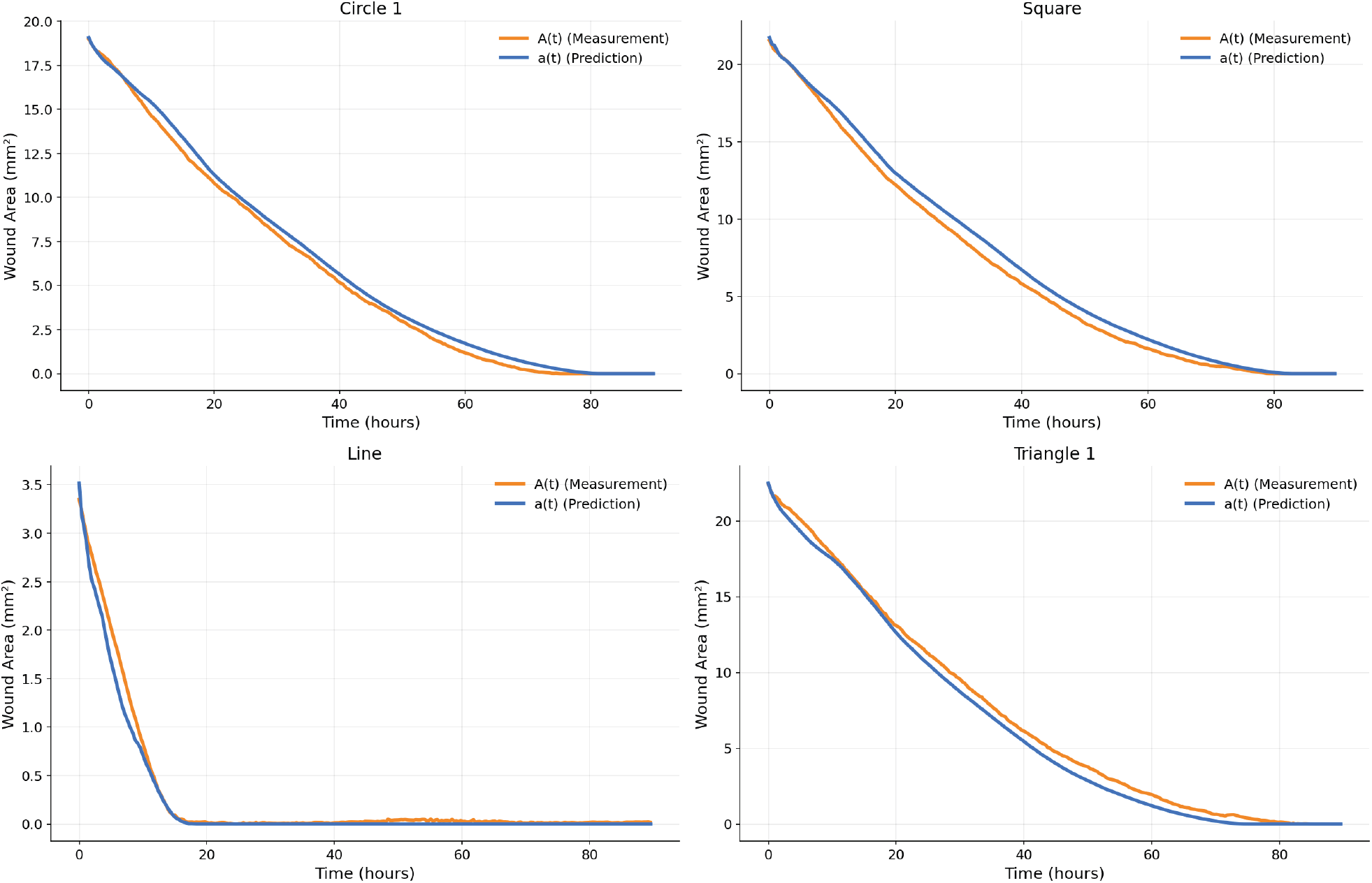
Representative wound area-time graph samples from each geometry type (Circle, Square, Line, Triangle) for the leave-one-out test.

Comparing Δ*T*_*c*_ between wound shapes, the circle datasets demonstrate positive Δ*T*_*c*_, indicating delayed closure relative to measurements (5h - 6h), while the triangles predict closure earlier than measured. The Triangle 3 dataset has the largest *n*Δ*T*_*c*_, and the Line dataset has the smallest.

### 3.2 Prediction Test

Figure 5 shows *a*(*t*) overlaid with *A*(*t*) over the last 66 hours for each of the Circle 1, Square, and Triangle 1 datasets after tuning the simulator using the first 24 hours. Table 3 lists NRMSE, *R*, Δ*T*_*c*_, and *n*Δ*T*_*c*_, after tuning the simulator using the first 24 hours of each video to predict wound closure in the remaining 66 hours. In this test, the Line dataset was omitted, as the wound in that dataset closes in less than 24 hours. The NRMSE stayed consistently low across all geometries (*µ* = 2.08%, *σ* = 2.29%), indicating a strong ability in predicting the wound area-time curve. The Pearson’s *R* remained *>* 0.98 for all cases, demonstrating strong agreement between *a*(*t*) and *A*(*t*) for the remaining 66 hours of each video. The mean *n*Δ*T*_*c*_ was 7.99%, with a sigma of 8.85%, showing a moderate agreement in closure times, albeit less strong than that of the leave-one-out test.

**Table 3:** Mean prediction model performance across datasets.

| Dataset | NRMSE | R | $\Delta T_c$ (h) | $n\Delta T_c$ |
| --- | --- | --- | --- | --- |
| Circle 1 | 1.10% | 0.9992 | 3.46 | 5.08% |
| Circle 2 | 1.24% | 0.9984 | -1.94 | 2.73% |
| Square | 0.57% | 0.9997 | -3.21 | 4.41% |
| Line | - | - | - | - |
| Triangle 1 | 1.65% | 0.9984 | -6.88 | 9.24% |
| Triangle 2 | 1.19% | 0.9995 | 0.98 | 1.30% |
| Triangle 3 | 6.70% | 0.9813 | -21.30 | 25.21% |
| $\mu$ | 2.08% | 0.9961 | -4.82 | 7.99% |
| $\sigma$ | 2.29% | 0.0073 | 8.82 | 8.85% |

**Figure 5:**
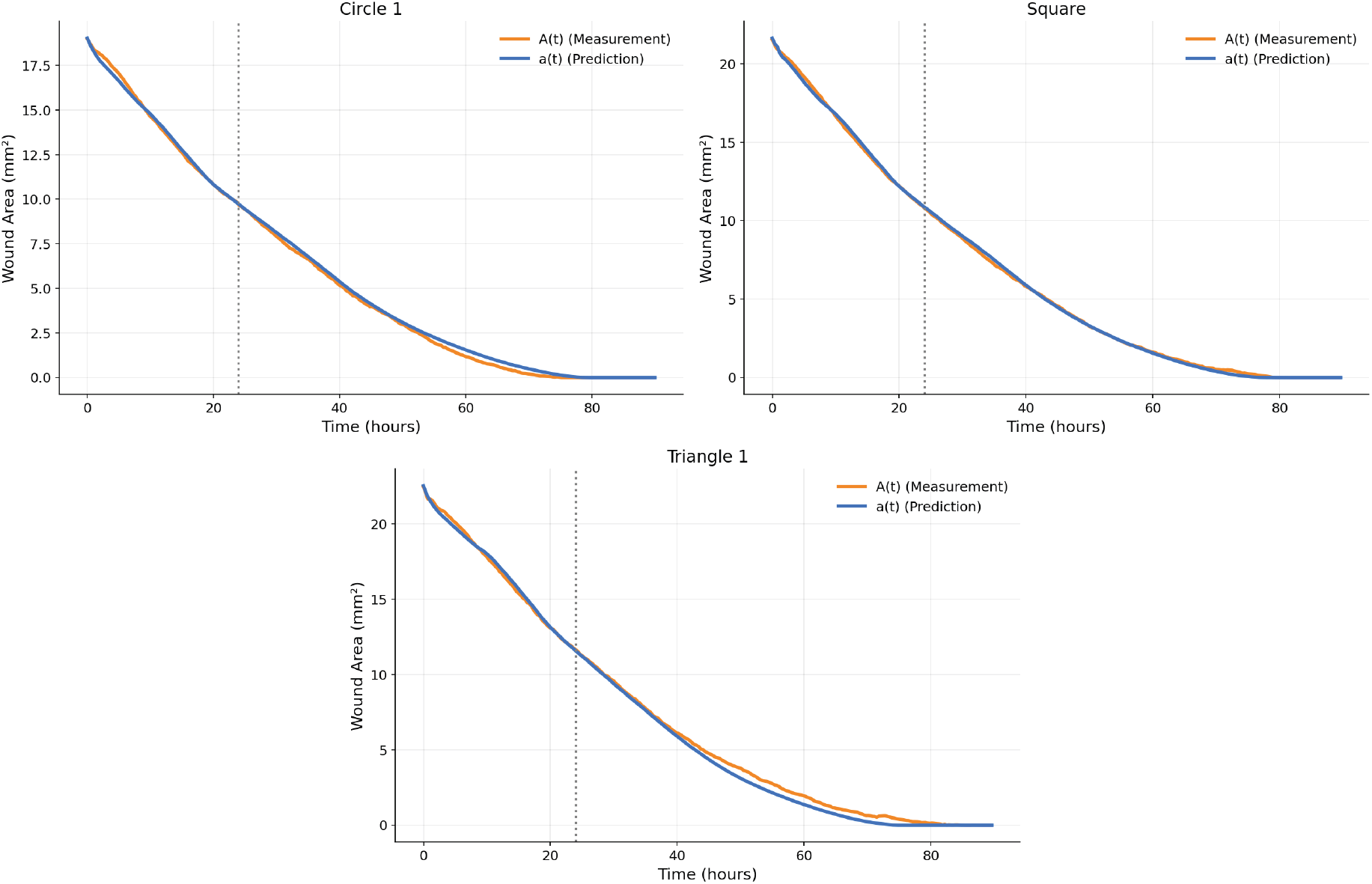
Representative wound area-time graph samples from each included geometry type (Circle, Square, Triangle) for the prediction test. The training and testing phases are separated by a vertical line at t=24.

Compared to the leave-one-out test, the prediction test achieved a mean NRMSE of 2.08%, while the leave-one-out test achieved a higher mean NRMSE of 3.34%. Between the two, the mean Pearson’s R remained consistent (0.9961 vs. 0.9989). However, the mean *n*Δ*T*_*c*_ increased, from 2.81% to 7.99%. In both tests, the Triangle 3 dataset performed the worst, with both the highest NRMSE and highest *n*Δ*T*_*c*_ across both datasets.

### 3.3 Sensitivity Analysis

Table 4 lists the results from the sensitivity analysis. Across all datasets, the *cell-repulsion* parameter consistently exhibits the highest *µ*^∗^, and parameter *ecm-speed-scalar* exhibits the second-highest, indicating that the model is most sensitive to these parameters. The turning-related parameters, *turn-speed* and *turn-chance* exhibit low *µ*^∗^ values in all selected datasets, indicating low sensitivity.

**Table 4:** Morris sensitivity analysis across select datasets.

| Parameter | Circle 1 |  | Square |  | Line |  | Triangle 1 |  |
| --- | --- | --- | --- | --- | --- | --- | --- | --- |
| | $\mu^*$ | $\sigma$ | $\mu^*$ | $\sigma$ | $\mu^*$ | $\sigma$ | $\mu^*$ | $\sigma$ |
| cell-repulsion | 1.742 | 2.033 | 2.120 | 2.325 | 0.611 | 0.653 | 2.370 | 2.520 |
| max-speed | 0.571 | 0.749 | 0.701 | 0.860 | 0.093 | 0.104 | 0.650 | 0.850 |
| turn-speed | 0.342 | 0.494 | 0.126 | 0.159 | 0.082 | 0.105 | 0.100 | 0.160 |
| turn-chance | 0.075 | 0.117 | 0.069 | 0.125 | 0.018 | 0.025 | 0.100 | 0.130 |
| inhibition-radius | 0.333 | 0.267 | 0.356 | 0.365 | 0.060 | 0.061 | 0.420 | 0.450 |
| ecm-avoidance | 0.035 | 0.047 | 0.023 | 0.030 | 0.049 | 0.108 | 0.680 | 1.360 |
| ecm-speed-scalar | 0.669 | 1.311 | 0.437 | 1.182 | 0.164 | 0.260 | 1.190 | 1.970 |

Comparing between datasets, the Triangle 1 dataset demonstrates higher *µ*^∗^ and *σ* values than the Circle 1 and Line datasets across over half of its parameters. The Line dataset, on the other hand, demonstrates lower *µ*^∗^ and *σ* values.

### 3.4 Spatial Analysis

Figures 6 and 7 provide a visual representation of the spatial dynamics in both the leave-one-out and prediction tests. Figure 6 shows the unsigned error (RMSE), while Figure 7 shows the signed Δ*T*_*c*_. The triangle datasets displayed unusual features, most prominently in Triangle 3, for both figures.

**Figure 6:**
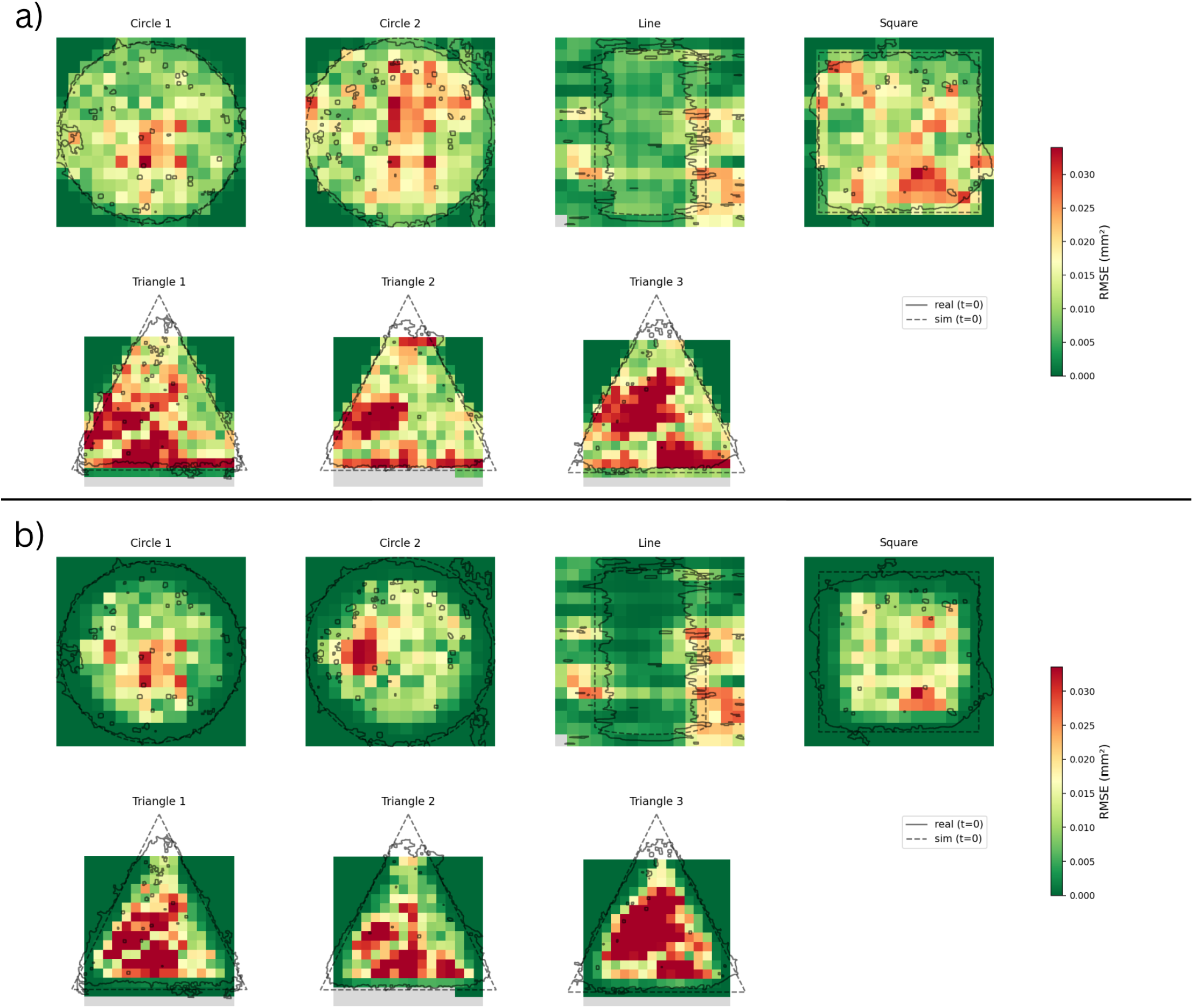
Heatmaps of the per-tile RMSE calculations for all datasets, in a) leave-one-out and b) prediction tests.

**Figure 7:**
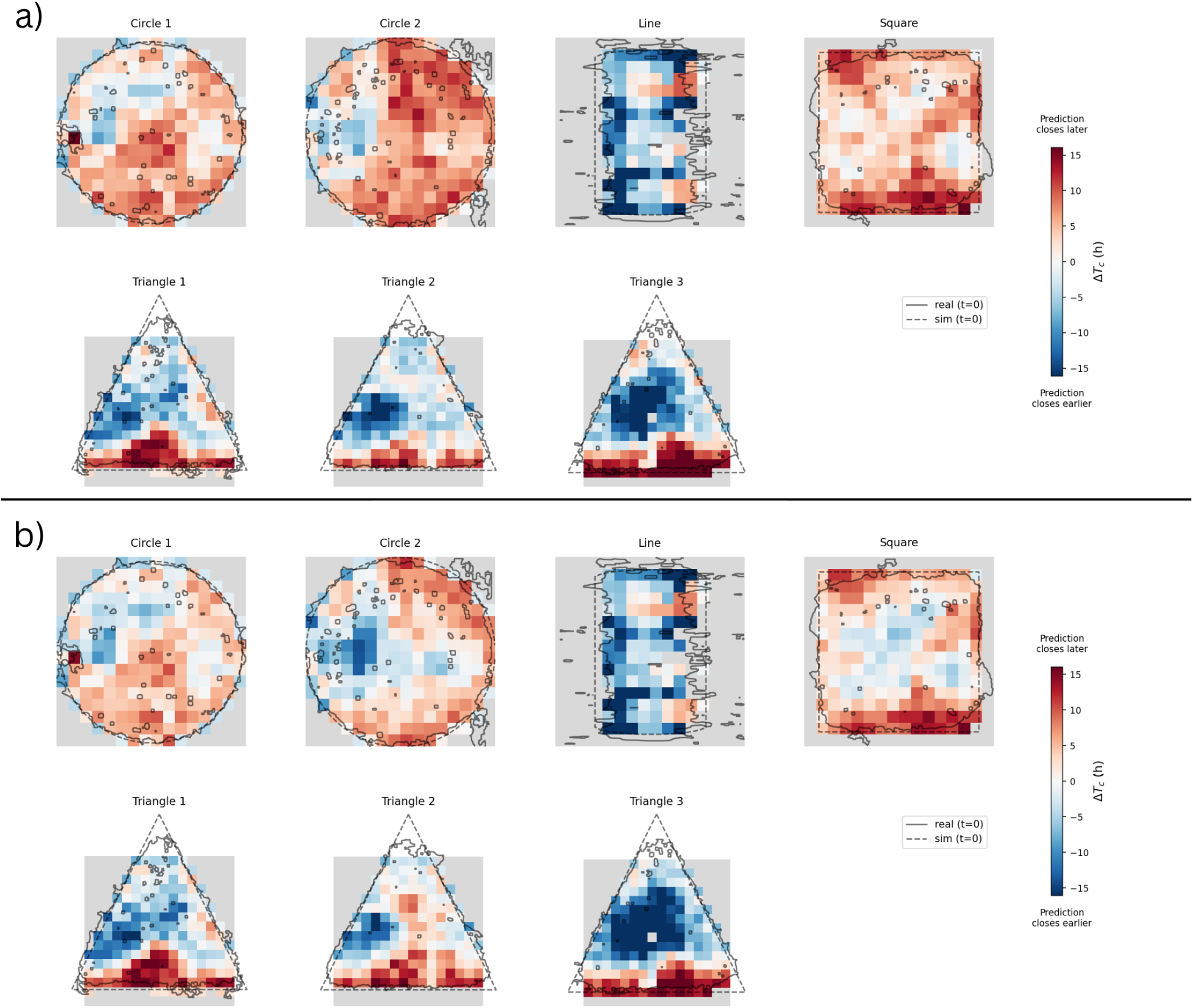
Heatmaps of the per-tile Δ*T*_*c*_ calculations for all datasets, in a) leave-one-out and b) prediction tests.

## 4 Discussion

The primary objective of this study was to determine whether wound closure dynamics observed experimentally can be modeled by a simulation with simple agent-based migration and proliferation rules. The results demonstrate that our model was able to accurately reproduce wound closure behavior across varying wound shapes, and that those behaviors were able to emerge from those agent-level rules. The high Pearson’s R (*>* 0.98 for each shape in both leave-one-out and prediction tests) and low error (NRMSE *<* 7% for each shape in all tests) indicate general agreement between experimental and simulated wound area-time curves.

Leave-one-out training was possible with the imaging dataset. Its success indicates that the model captures general characteristics of fibroblast migration and proliferation in wound healing rather than specializing to a single wound geometry. The model’s rules generalized to previously unseen wound geometries, indicating its robustness.

The ability of the model to predict broad wound area over time given only early closure data suggests that early dynamics may contain sufficient information to make assumptions about the subsequent closure behavior. The predictive capability of the simulation model suggests that with further validation, it may be used to predict wound closure times and increase the speed of experiments.

In the sensitivity analysis, the *cell-repulsion* simulation parameter consistently achieved the highest *µ*^∗^ (1.742, 2.120, 0.611, 2.370) across all tested datasets, indicating high impact on the model. This finding suggests that mechanical forces or heightened migration in response to rapid proliferation could play a large role in influencing the speed of wound healing [15]. Cell repulsion represents the tendency of cells to spread into regions of lower cell density to maintain even packing. The greater significance of cell repulsion compared to the motility parameters (i.e. *turn-speed*, and *turn-chance*) suggests that modeling the collective behavior of cell packing and density gradients may be more important than precisely modeling individual fibroblast migration patterns.

The *ecm-speed-scalar* parameter consistently achieved the second or third-highest *µ*^∗^ (0.669, 0.437, 0.164, 1.190). However, the other ECM-related parameter, *ecm-avoidance*, ranked significantly lower, although with more variability. In the Circle, Square, and Line datasets, it ranked among the lowest *µ*^∗^ values, while in the Triangle 1 dataset it held the third-highest *µ*^∗^. This indicates that changes in migration speed in response to ECM generally hold more significance in influencing wound healing than a directional change in our model. However, our representation of ECM as directionless particles instead of fibers may have influenced these results [8].

Our model is limited by a combination of factors in our design and validation procedures. Firstly, we include several processes that are not central to the validation procedure, such as the cell cycle and ECM deposition. The effects of these processes are not quantified or validated, meaning that we cannot conclude whether they had any significant effect on the model’s end goal. The use of area as the central validation method limits our conclusions to a global scope. Without a rigorous procedure that the spatiotemporal aspects of the simulation match with reality, we cannot conclude that the model replicates spatial wound closure dynamics with certainty. However, we can conclude that the model can predict the temporal dynamics of wound healing, such as predicting when a given wound will close. Our model is also limited by the size (7 shapes) and quality of the validation dataset. In the process of extracting the wound area from the imaging data, extraneous regions outside of the wound zone appeared due to the technology used to capture the images. In addition, errors in the stitching together of frames led to occasional sudden jumps in wound area. Both of these issues were mitigated, but only to an extent. The quantity and variety of geometries available also limited testing. In addition, the Triangle datasets, most prominently in Triangle 3, exhibited inconsistencies in wound closure dynamics when compared against the other datasets. Specific areas of the wound closed at slower rates during specific periods, altering the wound area time series. In order to determine whether this phenomenon was an error in data collection or a behavior of wound healing unaccounted for in our model, we would need further datasets to increase our sample size.

All datasets exhibited a significant jump in mean brightness and a horizontal shift at approximately t=71.8h. This disruption was caused by restarting the imaging microscope, and a simultaneous media change. To mitigate the effects of the disruption, we corrected the wound area calculations by setting the area following the jump equal to the area prior to the jump. The disruption could be a factor in the relatively large error in some datasets past t=71.8h. We observed an approximately linear streak pointed towards the center of the wound area in the Triangle 3 dataset, and hypothesized that the streak could be a factor in the large error in the Triangle 3 dataset across all tests. However, the streak could have been created in the imaging process rather than in fibroblast behavior. We hypothesized that the streak was likely either an illumination artifact that was corrected once the camera or LEDs were reset during the restart, or condensation on the plate/lid. During data collection, restarting the microscope usually ejected the plate from the incubation chamber, so the brief temperature and humidity change could have cleared or redistributed the condensation.

In future work, we plan to improve our validation procedure, further investigate the effects of wound curvature and spatial features on wound closure, and extend the model towards clinical relevance. In order to improve the robustness of our validation method, we plan to use edge-detected contour shapes as the initial conditions for wound boundaries as opposed to ideal geometric approximations, test our simulation against further scratch assay data from different sources and under different experimental conditions, and expand upon and validate some of the unvalidated processes in the model such as the cell cycle and ECM deposition. In addition, we intend to analyze the effects of wound boundary curvature, and the effects of different shapes on the wound area-time graph. To extend the significance of our model, we aim to validate our model in more complex in vitro scenarios, such as multi-layered scratch assays, or mixed assays with keratinocytes. Then, we hope to extend the principles of the model to in vivo scenarios.

In this paper, we developed an agent-based model of fibroblast migration and proliferation in wound healing scratch assays that simulates wound closure using simple local rules governing cell motility, contact inhibition, interaction with the extracellular matrix, and mechanical forces. Our model reproduced experimentally observed wound closure dynamics through both a leave-one-out test and a prediction test, demonstrating the model’s ability to generalize across wound geometries, and its ability to predict wound closure given only the first 24 hours of data. These results demonstrate that high-level wound closure behavior can emerge from a relatively small set of local, agent-based rules. As a computationally efficient and interpretable framework, our model provides a platform for investigating and predicting wound healing dynamics in scratch assays.

## 5 Acknowledgments

This work was supported by the use of generative AI agents including Claude and ChatGPT. Generative AI was used for background research, coding, and editing. All AI-generated work was reviewed and fact-checked by human authors.

## 6 Code Availability

The simulation source code and analysis scripts are available at https://github.com/rowansuma/FISS.

## A Extra Figures

### A.1 Leave-one-out Test

Figures 8-14 display our optimized wound area-time graphs from the leave-one-out test.

**Figure 8:**
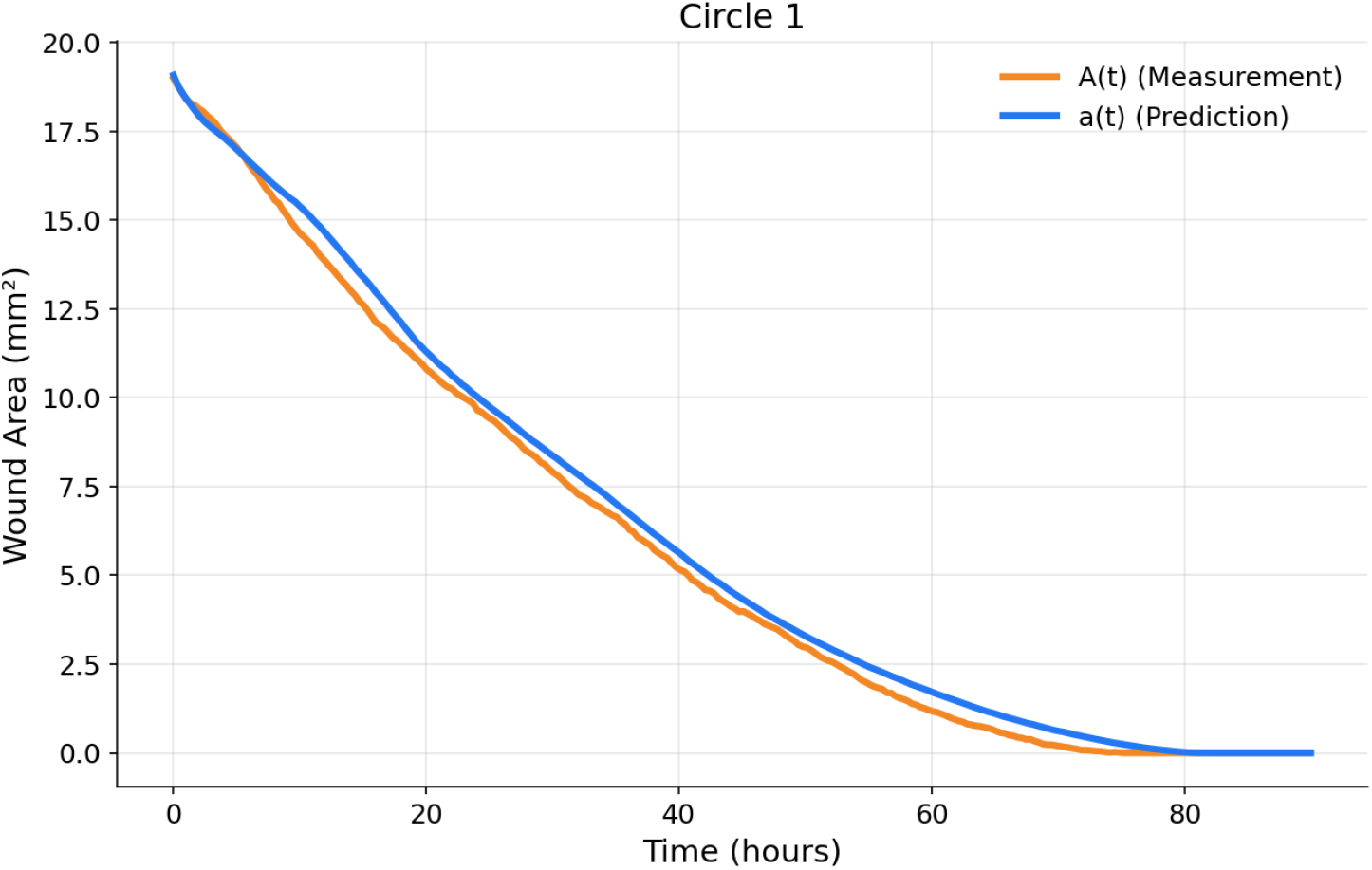
Leave-one-out test wound area-time graph for the Circle 1 dataset.

**Figure 9:**
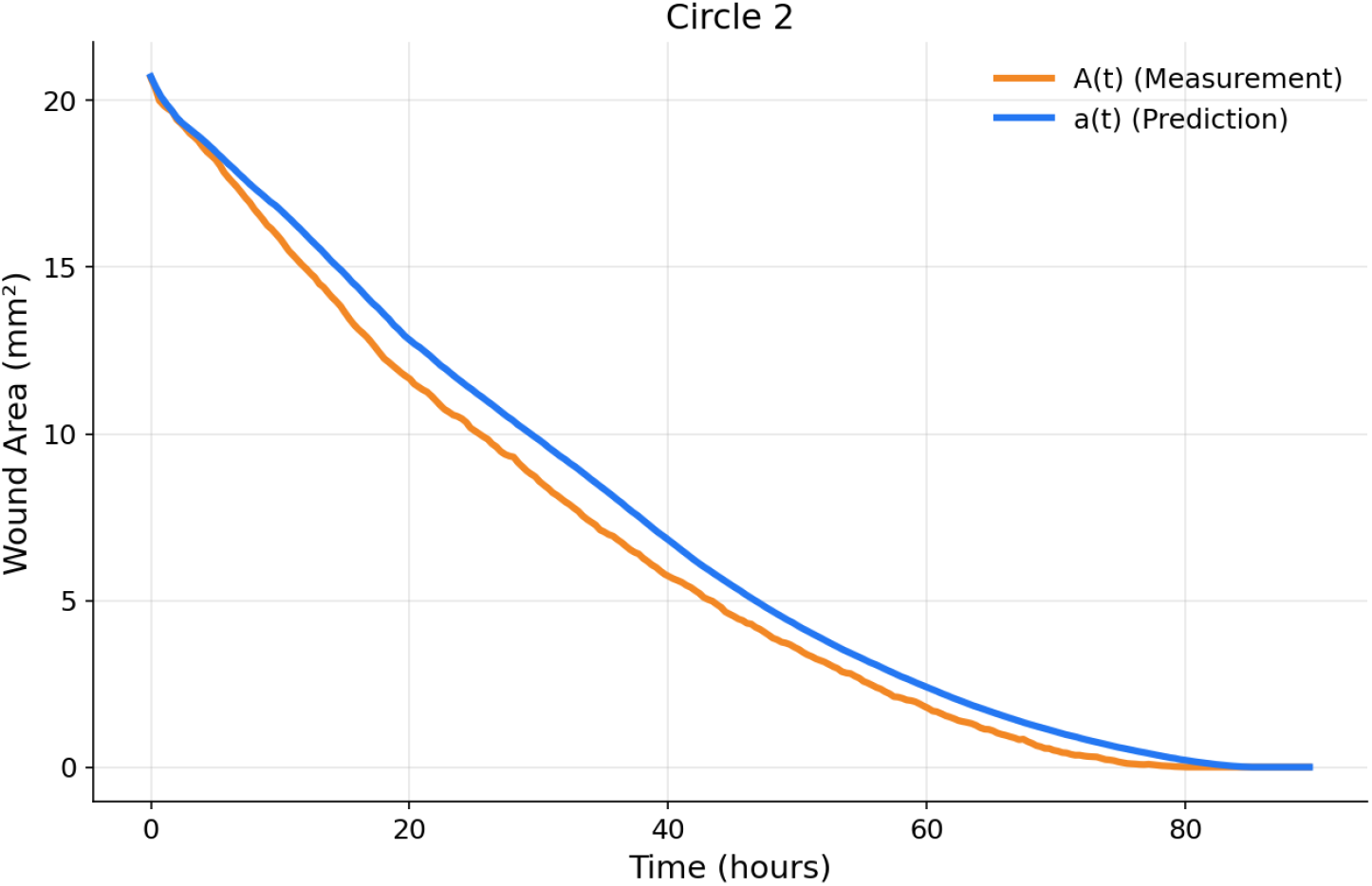
Leave-one-out test wound area-time graph for the Circle 2 dataset.

**Figure 10:**
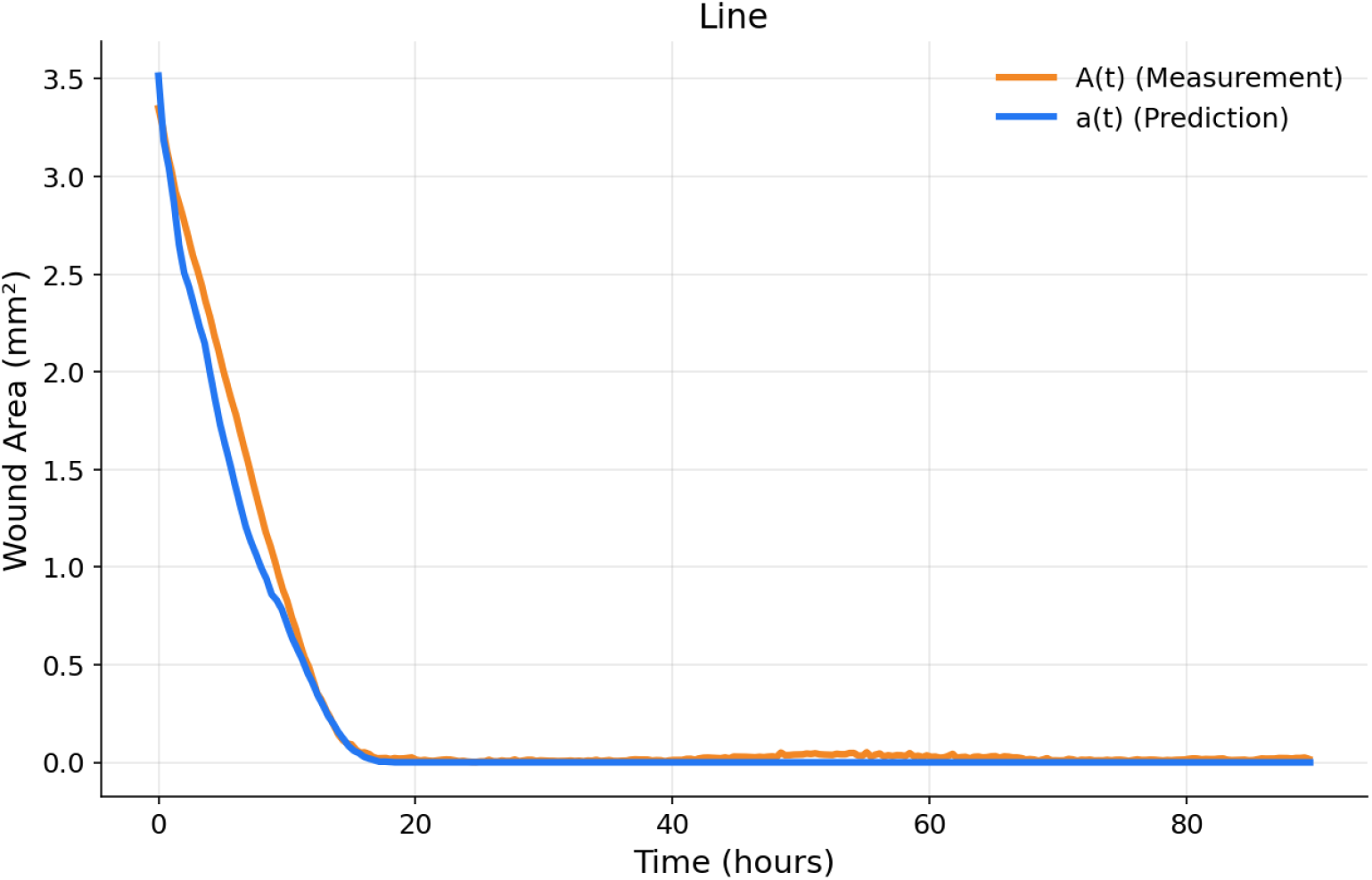
Leave-one-out test wound area-time graph for the Line dataset.

**Figure 11:**
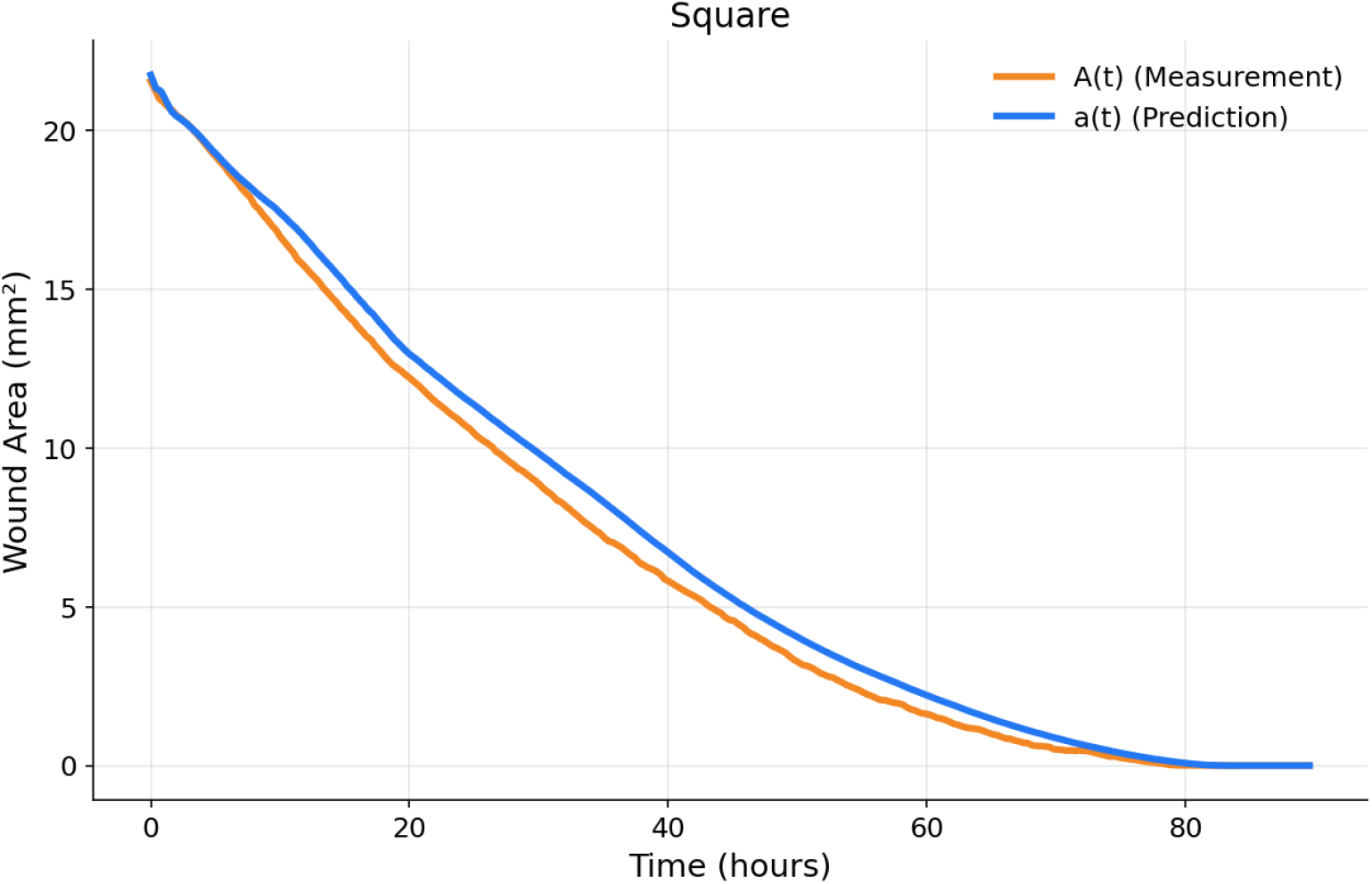
Leave-one-out test wound area-time graph for the Square dataset.

**Figure 12:**
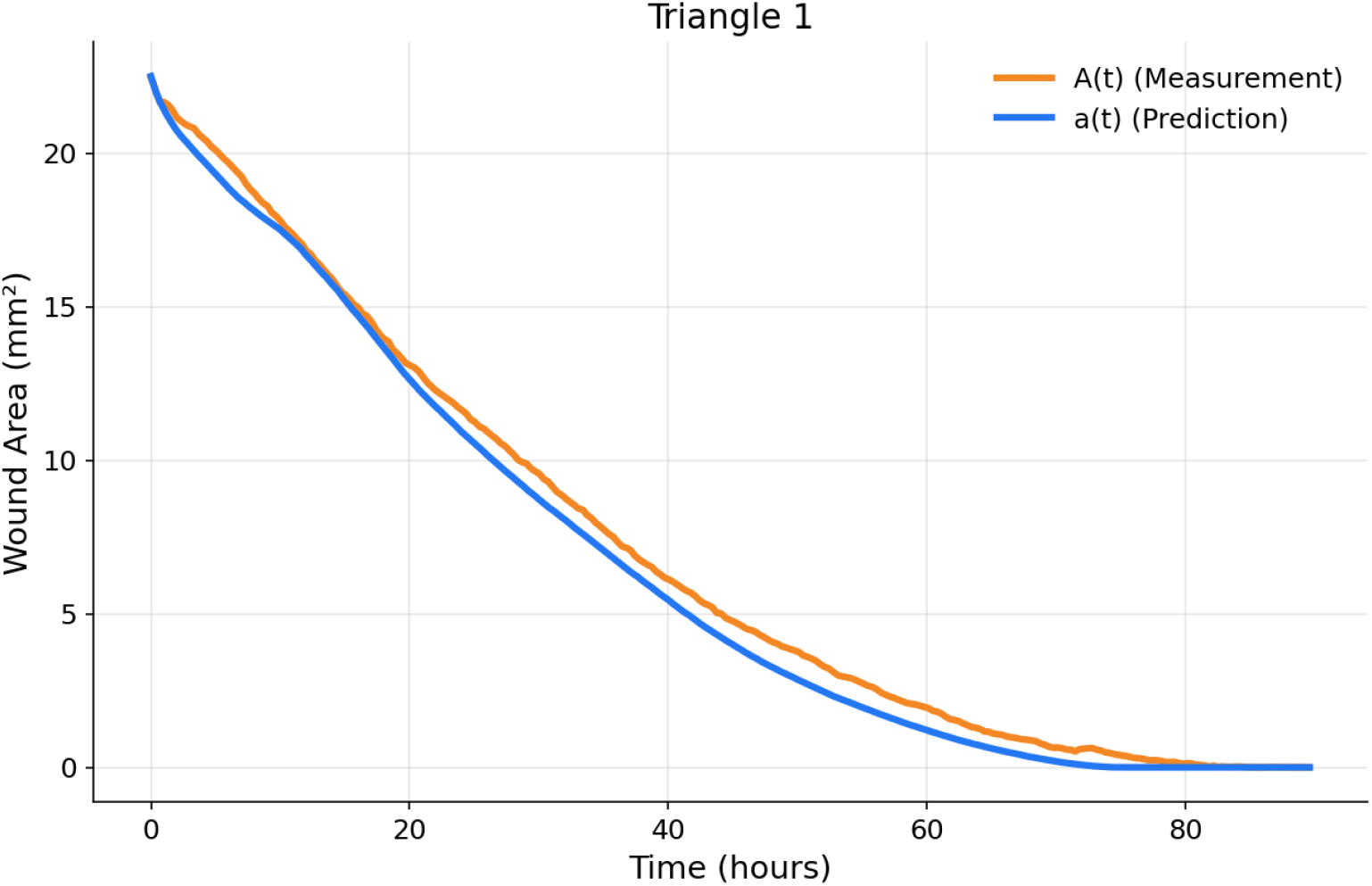
Leave-one-out test wound area-time graph for the Triangle 1 dataset.

**Figure 13:**
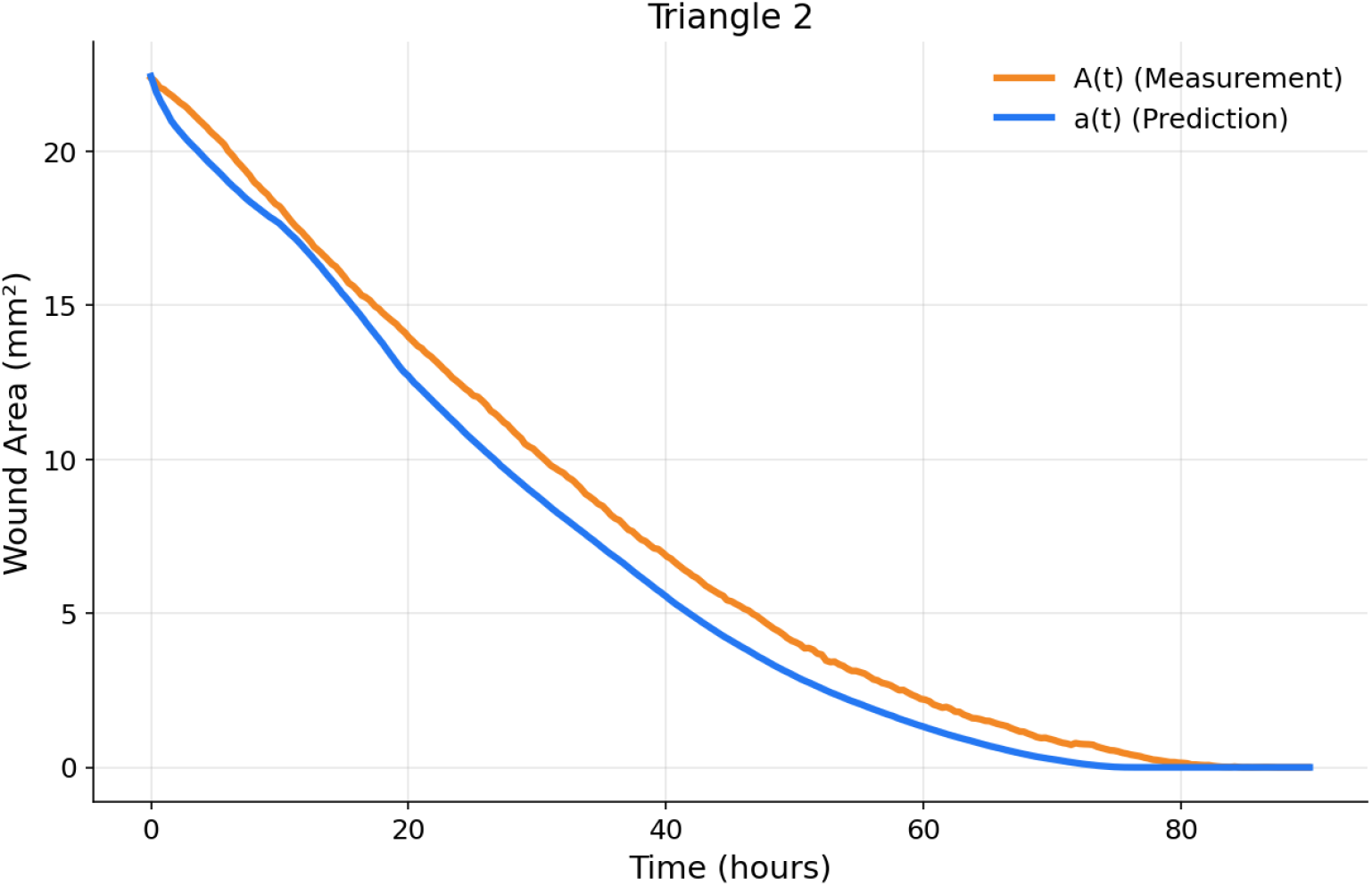
Leave-one-out test wound area-time graph for the Triangle 2 dataset.

**Figure 14:**
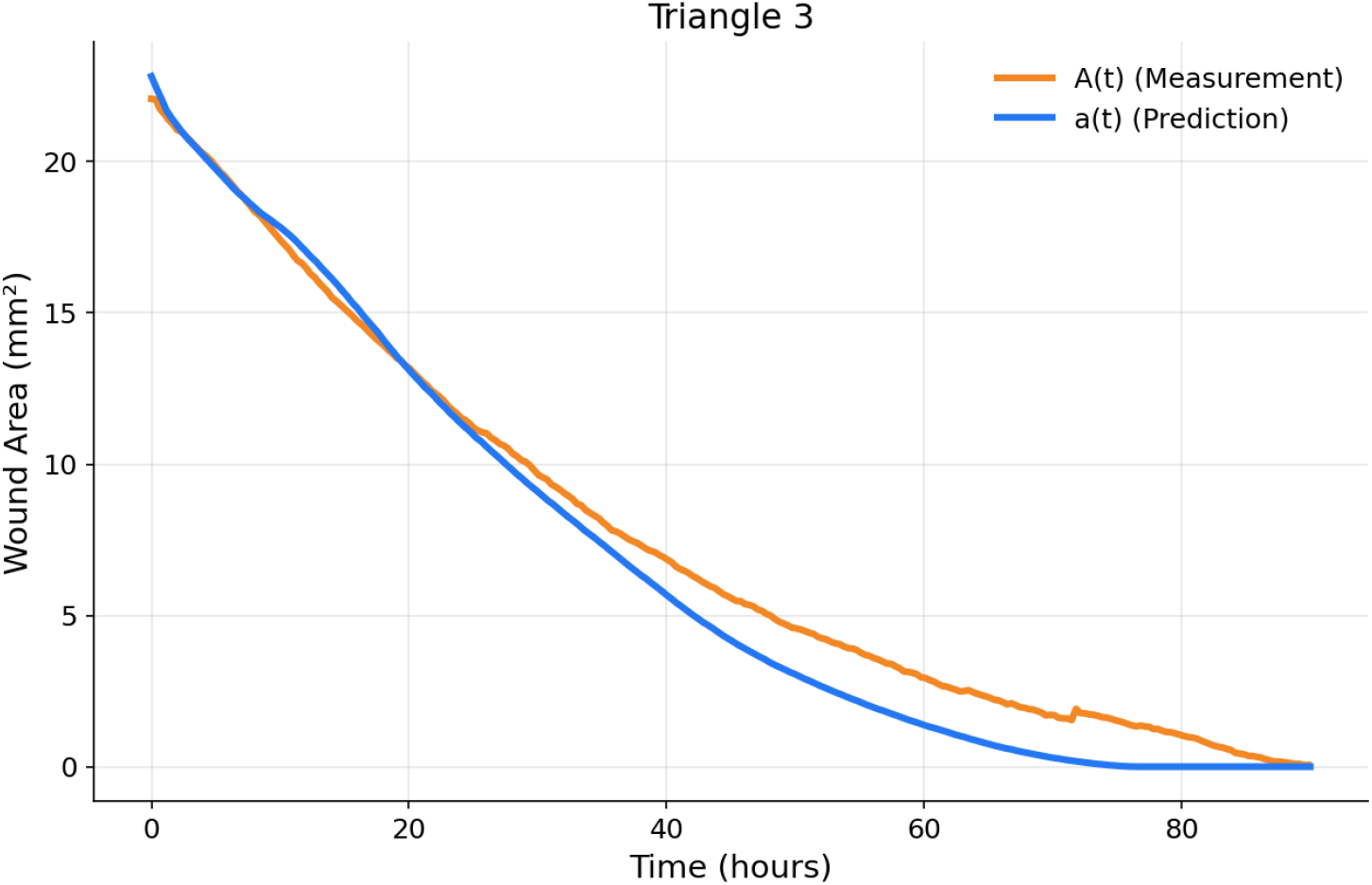
Leave-one-out test wound area-time graph for the Triangle 3 dataset.

### A.2 Prediction Test

Figures 15-20 display our optimized wound area-time graphs from the prediction test.

**Figure 15:**
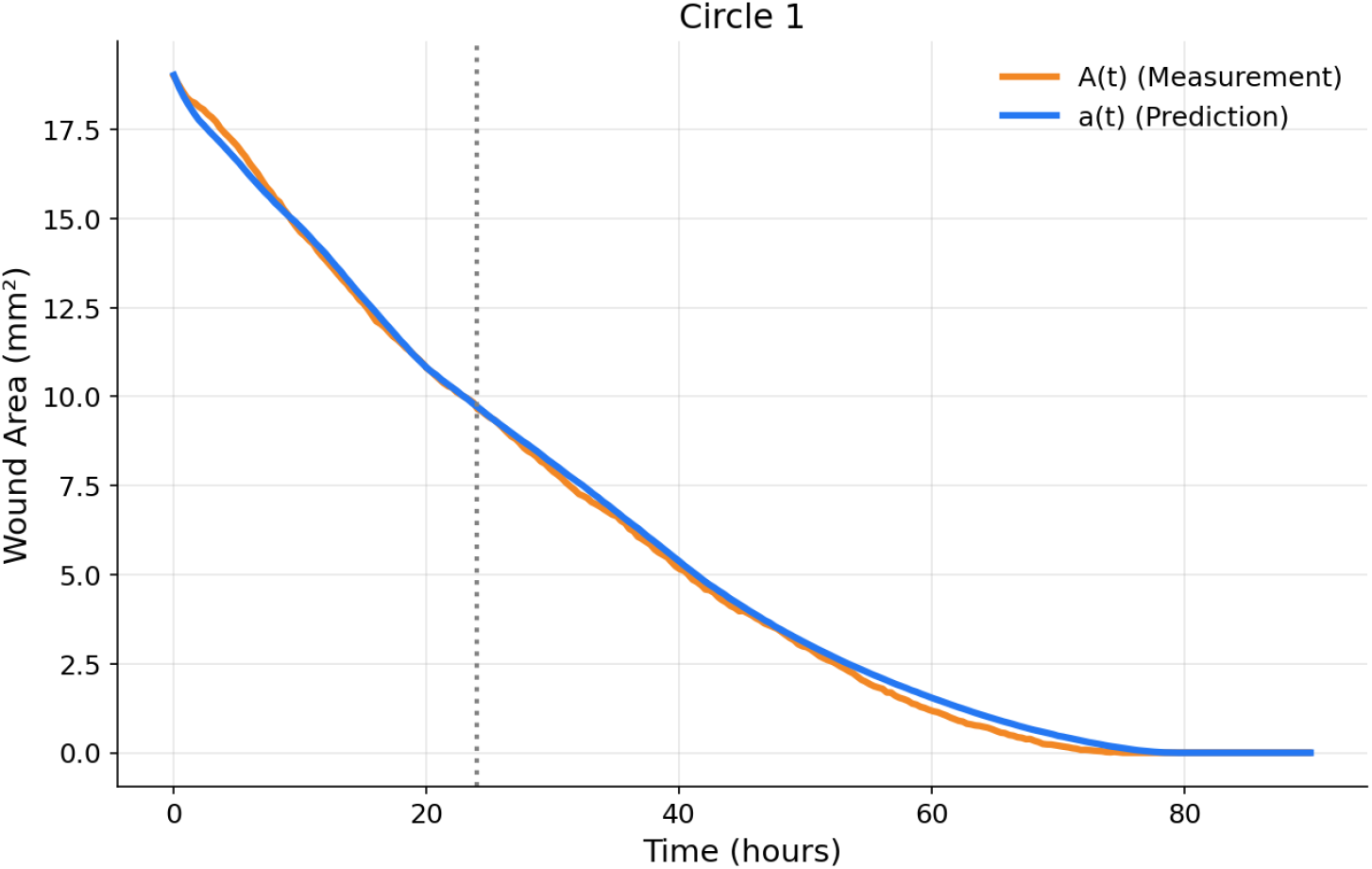
Prediction test wound area-time graph for the Circle 1 dataset.

**Figure 16:**
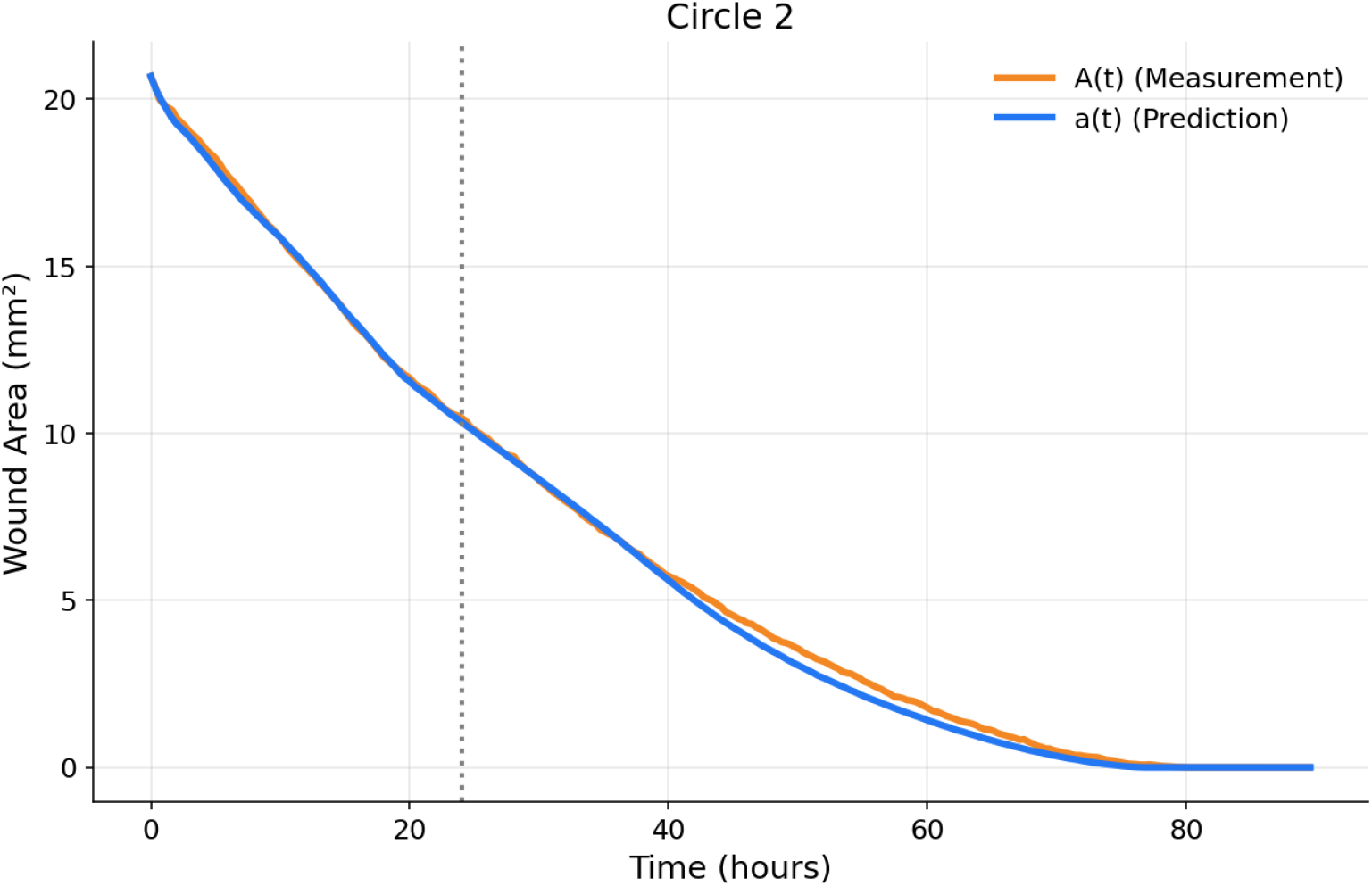
Prediction test wound area-time graph for the Circle 2 dataset.

**Figure 17:**
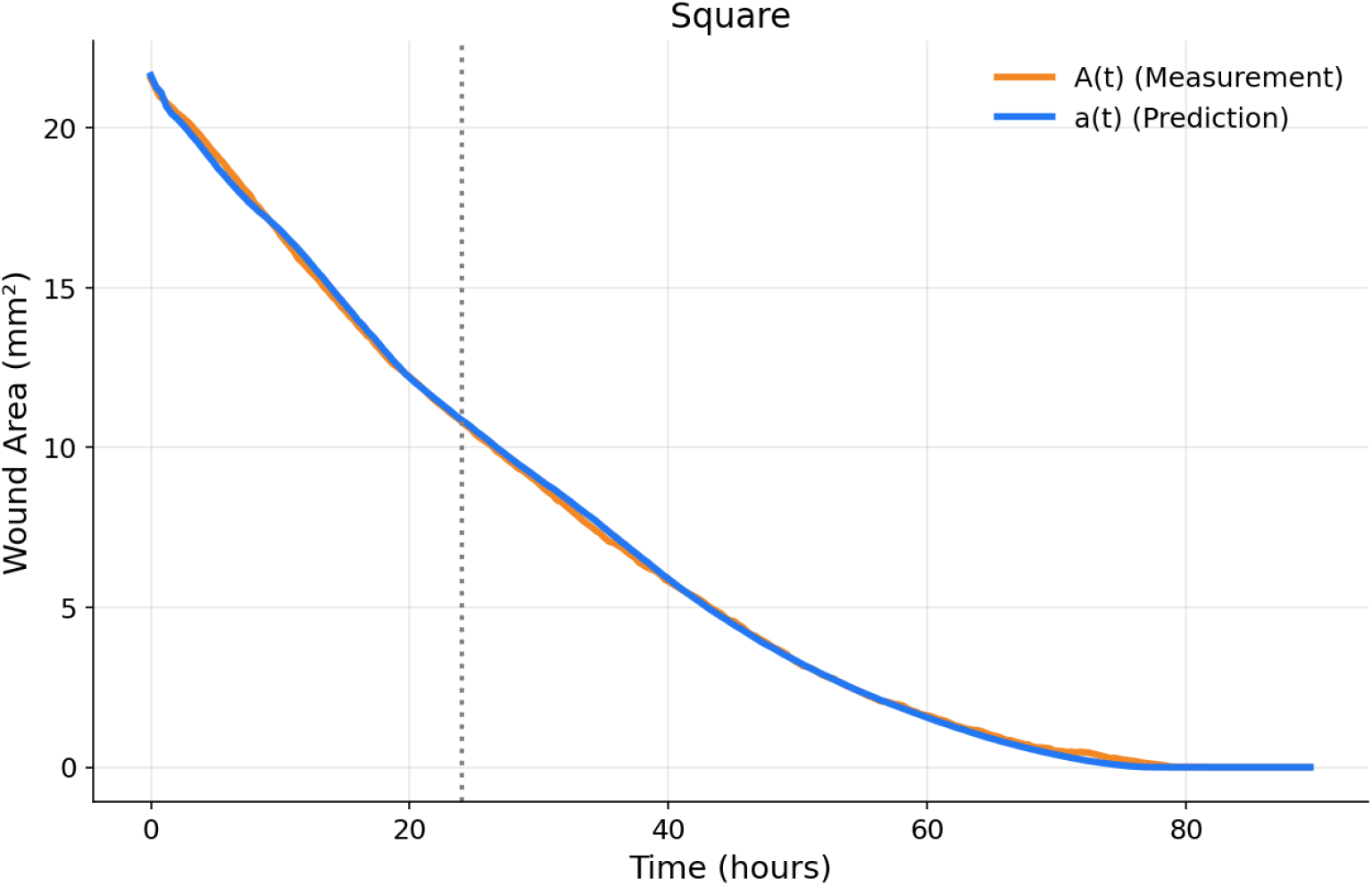
Prediction test wound area-time graph for the Square dataset.

**Figure 18:**
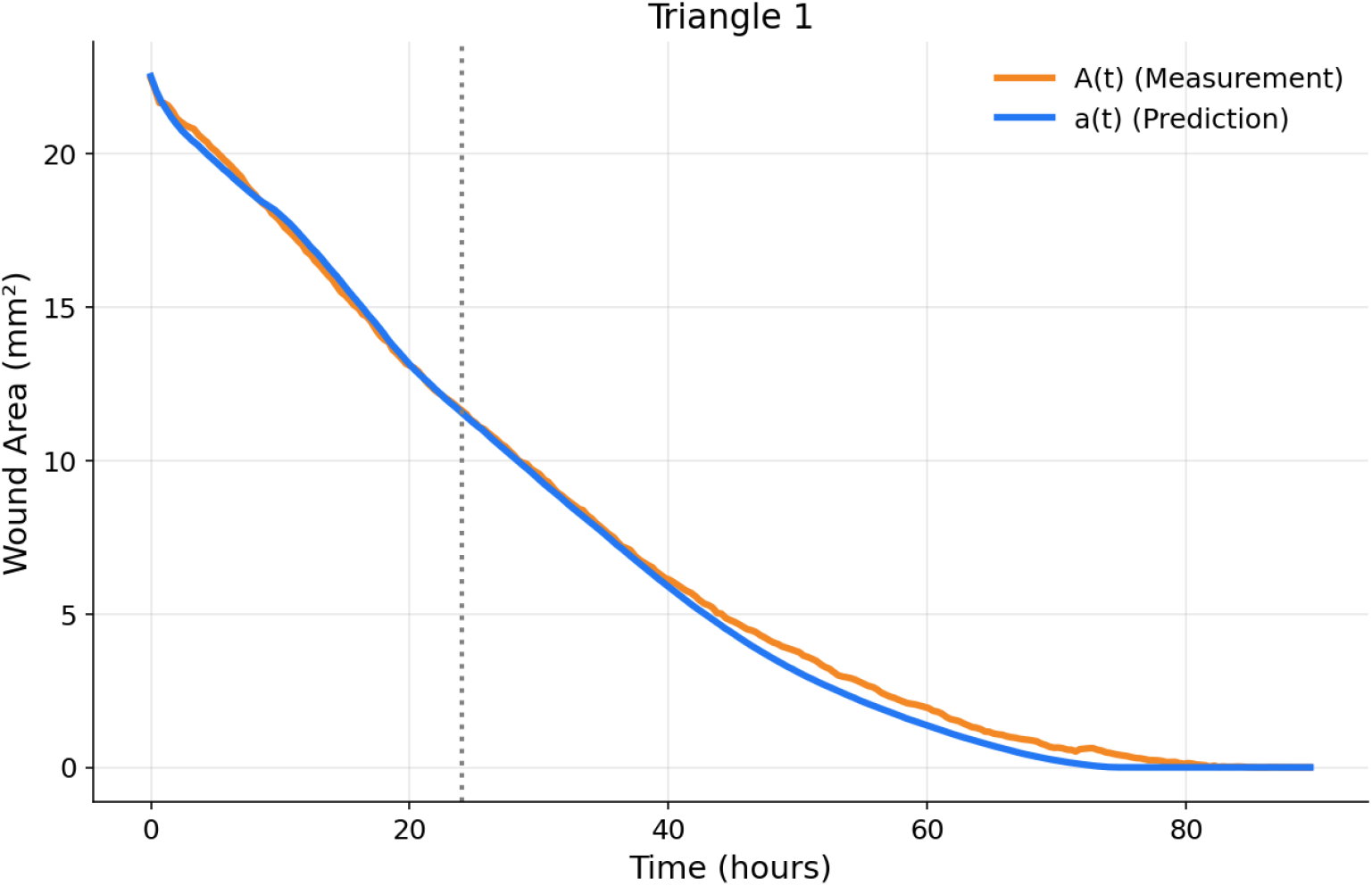
Prediction test wound area-time graph for the Triangle 1 dataset.

**Figure 19:**
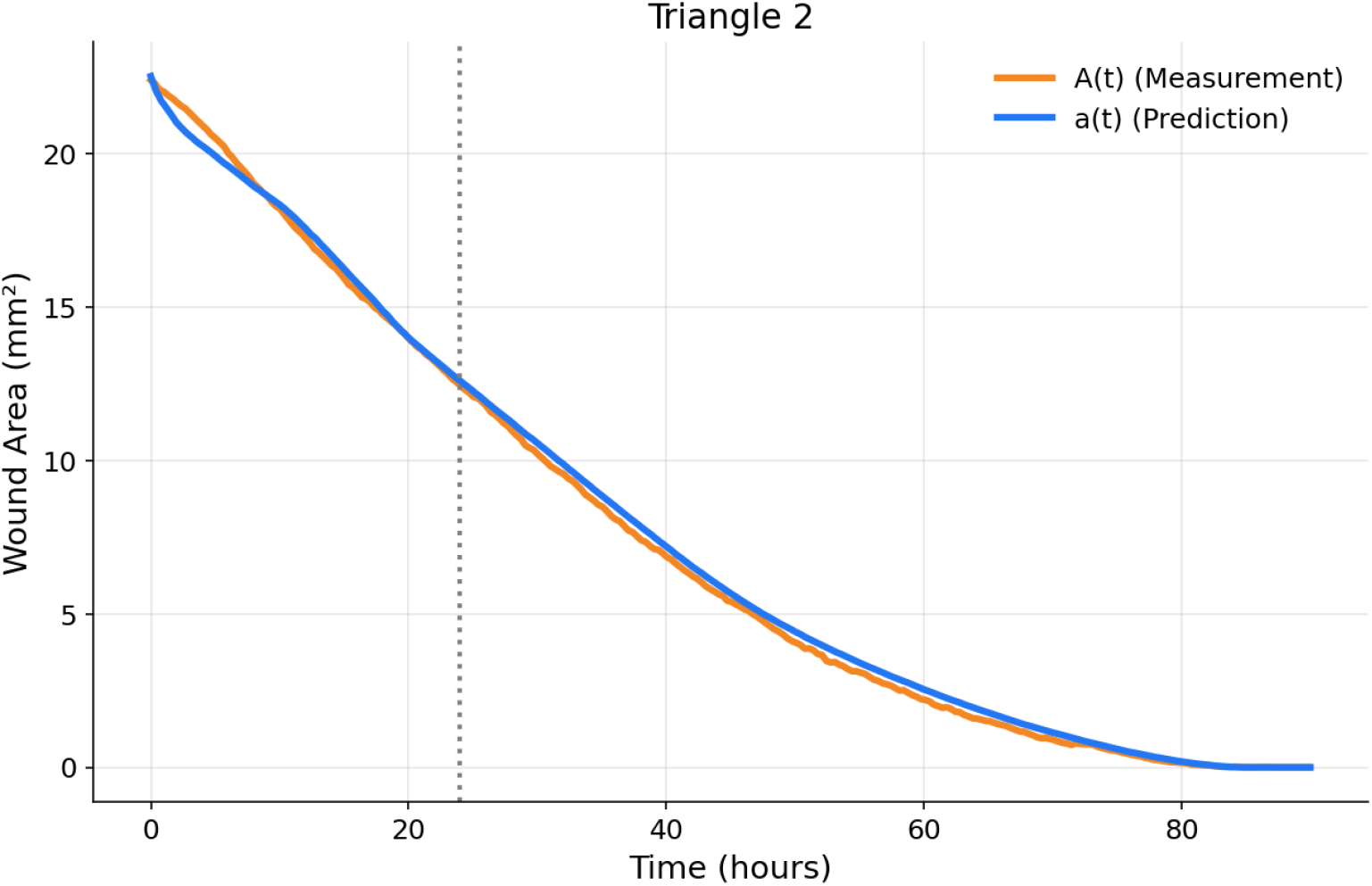
Prediction test wound area-time graph for the Triangle 2 dataset.

**Figure 20:**
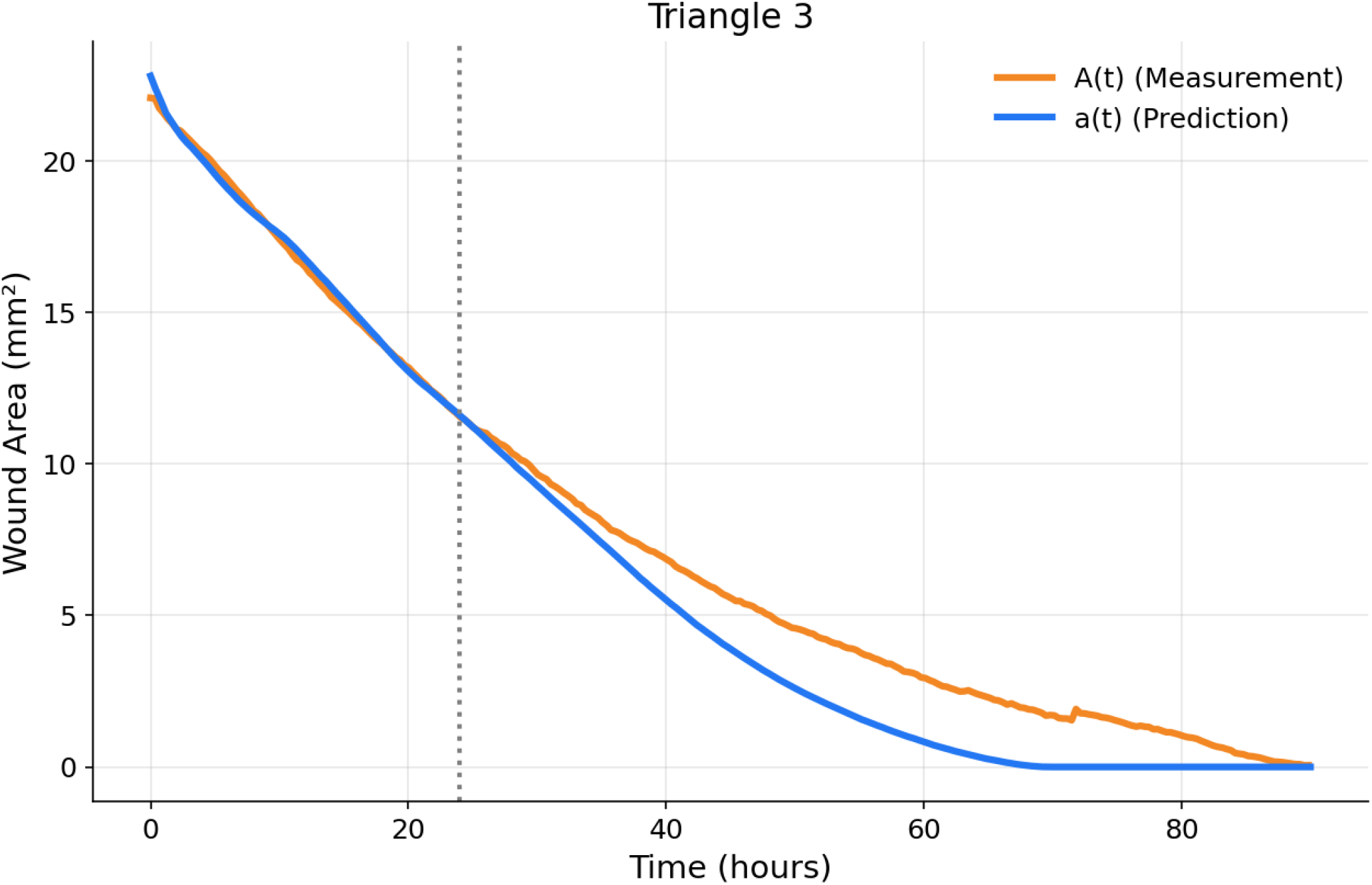
Prediction test wound area-time graph for the Triangle 3 dataset.

## B ODD Protocol

In this appendix, we describe our model according to ODD [10, 11, 12]. We follow “S1: ODD Guidance and Checklists” for guidance in writing ODD descriptions of ABMs, based on the ODD version published in [12].

### B.1 Overview

#### B.1.1 Purpose and Patterns

The immediate purpose of the model is to predict the dynamics of fibroblast proliferation and migration in a scratch assay monolayer across different wound configurations. The ultimate purpose of the model is to reduce the cost, time, and reproducibility problems conventional scratch assays face.

We evaluate our model by its ability to reproduce two patterns found in our test data.

*Pattern 1. Wound area reduction over time*. In scratch assay experiments, wound area reduces monotonically over time as cells proliferate and migrate into the wound region. The model aims to reproduce the different wound area-time curves observed in the test data across different wound geometries.

*Pattern 2. Healing times*. Many experimental scratch assays use the time required for a wound to reach a specific closure fraction (50%, 80% closed) as a benchmark of closure speed. The model aims to reproduce the experimentally observed time to 98% closure in our test data.

#### B.1.2 Entities, State Variables, and Scales

##### Entities

The model is composed of the following entities: agents representing fibroblast cells, particles representing the extracellular matrix (ECM), and the global environment representing the petri dish used in traditional scratch assays. ECM particles are static after deposition and do not decay or move.

##### State Variables

In Table A1 we show the entities’ state variables.

##### Scales

Simulation space dimensions were variable between different wound configurations because test data dimensions varied. Simulation step length is adjustable but is fixed to 2.5 hours per step for the purposes of this publication. All time-related parameters (e.g., cycle duration) are expressed in simulation steps.

#### B.1.3 Process Overview and Scheduling

The model proceeds in discrete timesteps. Each agent executes the following processes in sequence. Agents are updated in an arbitrary order.

1. Physics. Each fibroblast agent executes the “physics” submodel, which updates the agent’s position based on its previous position and applies physical constraints such as collision handling and boundary conditions.
2. Locomotion. Each fibroblast agent executes the “locomotion” submodel. The agents’ turn-state variable is updated stochastically, and their position is updated based on local ECM density and their current speed and turn-state variables.
3. Cell cycle. Each fibroblast agent executes the “cell cycle” submodel, which advances the agent’s cycle phase according to a configured timer. When a cell exits the M phase, a new fibroblast agent is created at the parent cell’s position.
4. ECM deposition. Each fibroblast agent executes the “ECM deposition” submodel, which deposits an ECM particle at the agent’s position according to a configured timer.
5. The display and file outputs are updated.

**Table A1:**
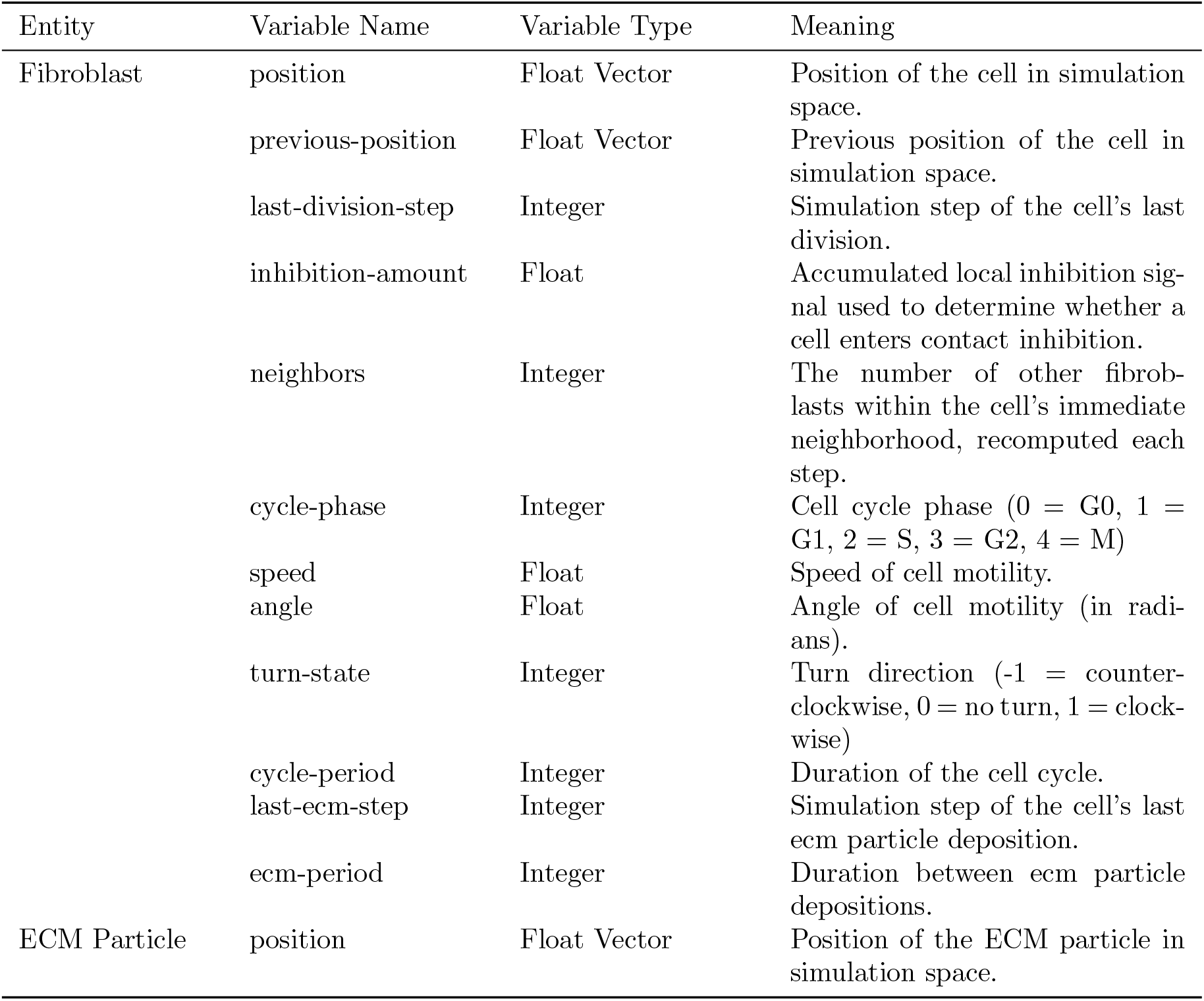
Entity state variables.

### B.2 Design Concepts

#### B.2.1 Basic Principles

The model follows an agent-based modeling approach that represents each fibroblast explicitly as a circle, in contrast with a continuum-based [1] or Cellular Potts [9] model. The model is based on four fundamental simulation and fibroblast-specific agent-level principles based on existing theory.

- Stochastic motility: cell motility is based on both a stochastically determined movement vector and environmental cues such as the local density of ECM particles and other cells.
- Mechanical exclusion due to physical crowding: cells exert repulsive forces on nearby cells to prevent overlap, representing crowding effects within confluent monolayers, and governing system-level behavior in confluent regions.
- Contact inhibition: cell proliferation and G0 exit/entry depends on local crowding.
- Phenotypic plasticity in response to ECM: high concentrations of ECM slow down migration and further deposition, while low concentrations speed up migration and direct the cell down the concentration gradient.

The model applies these principles through the physics, locomotion, cell cycle, and ECM deposition submodels. System-level dynamics such as confluent monolayer sheet formation and wound closure emerge as a result of these agent-level processes. At this stage, the model does not aim to provide insights about these basic principles, but instead reproduces experimentally observed dynamics in scratch assays.

#### B.2.2 Emergence

The key outcomes of the model are wound closure dynamics, healing patterns dependent on wound geometry, confluent monolayer formation, and differences in local cell density between the edge and center of a wound. These outcomes are not imposed at the system level, and emerge from agent-level processes (migration, proliferation), cell-cell interactions (physical repulsion, contact inhibition), and ECM-cell interactions (ECM avoidance).

#### B.2.3 Adaptation

Cells have three adaptive behaviors: deciding whether or not to enter quiescence, altering their rates of ECM deposition and locomotion in the presence of high ECM concentration, and directing locomotion based on local ECM concentration. These decisions are modeled as indirect objective seeking through the following conditions:

1. A cell enters quiescence if the number of neighbor cells in the local neighborhood exceeds a certain threshold.
2. A cell’s rate of ECM deposition and movement speed decreases proportional to the local concentration of ECM particles.
3. A cell’s movement is biased by the local ECM gradient, with preferential migration toward regions of lower ECM concentration.

#### B.2.4 Objectives

Objectives are not implemented in the current model. Agents have no explicit goals.

#### B.2.5 Learning

Learning is not implemented in the current model. Agents do not change their adaptive behavior over time.

#### B.2.6 Prediction

Agent-level prediction is not implemented in the current model. Agents do not predict future environmental states.

#### B.2.7 Sensing

Cells are assumed to perfectly sense the number of neighboring cells in direct contact, representing cell-cell mechanotransduction [15] processes that govern contact inhibition. Cells also sense the local concentration of ECM particles within a specified radius, representing cell-ECM mechanosensing [2] that governs migration. This radius is constant over time and among cells, and its value is ECM Deposition submodel parameter *ecm-detection-radius*.

#### B.2.8 Interaction

Cells interact directly with others in their local neighborhood through both mechanical repulsion and contact inhibition. Cells exert repulsion forces on any neighboring cells in direct contact to minimize overlap, and undergo contact inhibition based on its number of neighbors. Mediated interactions between cells occur through the shared medium of ECM particles by depositing in response to low ECM concentration, and reducing migration and deposition in high concentrations. These interaction mechanisms reflect cell-cell mechanical interactions and cell-matrix mechanosensing interactions observed in fibroblast monolayers.

#### B.2.9 Stochasticity

Stochasticity is used in two ways. First, upon the initialization of a cell, some state variables are stochastically assigned, specifically position, cycle-period, and ecm-period. Second, cell motility is partially dictated by stochastic factors; the state variable *turn-state*, which controls movement direction, is randomly updated at random intervals. This process is further detailed in Section 3.3.2.

#### B.2.10 Collectives

No explicit collectives are implemented in this model. However, cells collectively form a monolayer as cells proliferate. This process is entirely emergent; cells form a monolayer as a result of simple physical repulsion and contact inhibition rules. No state variables are assigned to this structure.

#### B.2.11 Observation

Real time visual output on the model interface displays the location of each cell and its cell cycle phase, via colored circles. ECM particles are also represented as smaller circles. Summary statistics on the cell population, ECM population, and wound area are also provided through plots on the interface and output files. If enabled, imaging output designed to approximate microscopic imaging, and individual cell paths can also be recorded. All summary statistics are recorded every 30 simulation steps.

### B.3 Details

#### B.3.1 Initialization

In order to initialize a wound healing simulated experiment, the model requires the pre-generation of a confluent monolayer beforehand. This can be implemented by initializing one or more cells and allowing the simulation to run until all space is filled with quiescent cells. Then a wound can be created, and the experiment can begin. This method simulates real-world culturing of fibroblasts before a scratch assay, and randomizes spatial organization of cells in the final monolayer. Through this process, the position, last-division-step, neighbors, cycle-period, last-ecm-step, and ecm-period state variables for each fibroblast are stochastically determined without explicit initialization. At the start of the experiment, the system consists of a population of fibroblast cells and a population of ECM particles filling the entire domain, with a vacant area in the center of the domain representing the wound. The wound shape is dictated by the wound-shape parameter, which can be configured to be a circle, triangle, square, or line.

#### B.3.2 Input Data

The model does not use input data to represent time-varying processes.

#### B.3.3 Submodels

##### Physics

Mechanical interactions between cells are modeled by a Position-Based Dynamics (PBD) [14] approach. In implementation, positional adjustments are applied directly to state variable *position* during a simulation step. For clarity of presentation, that process is expressed here as a summation of incremental displacements at the end of a timestep.

Cells are represented as circular particles with a radius defined by submodel parameter *cell-radius*. In order to minimize overlap, cells are repulsed according to the following equations.

Let each cell have the following properties:

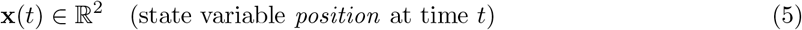

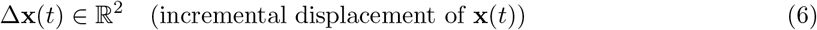

Each step, Δ**x**(*t*) is defined by

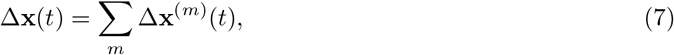

where *m* ∈ {phys, loc, ecm}. Changes in position in each submodel are accumulated in Δ**x**(*t*), which is updated at the end of each step by

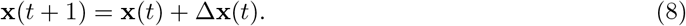

First, the submodel handles cell-cell repulsion. For each cell *j* and each other cell *k*, let the distance between their centers be

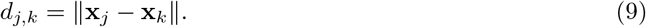

If the cells overlap, i.e., 0 *< d*_*j,k*_ *<* 2*ρ*, where *ρ* is submodel parameter *cell-radius*, then a cell-cell repulsion vector is applied:

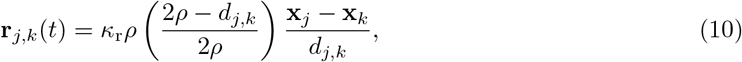

where *κ*_r_ is the submodel parameter *cell-repulsion*. This vector points away from the cell *k* and has a magnitude proportional to the overlap distance (2*ρ* −*d*_*i,j*_).

The physics submodel is run several times to ensure minimal collision. Model parameter *substeps* is the number of times the submodel is run. Changes to position are accumulated in Δ**x**^(*phys*)^(*t*), and*\* the substep iteration is defined by

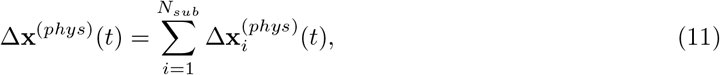

where *i* is the current substep and *N*_*sub*_ is submodel parameter *substeps*.

Cell positions are updated each substep according to

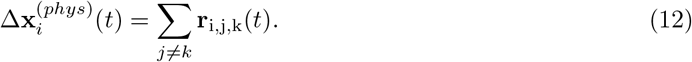

Boundary constraints are then applied by snapping cell positions to the edge of the simulation space if they are out-of-bounds.

After this process, state variable *inhibition-amount* is incremented for every neighboring cell within a fixed radius defined by the cell cycle submodel parameter *inhibition-radius*. This step is included in the Physics submodel to reduce the number of redundant neighbor iterations and increase efficiency.

In Table A2 we show the physics submodel parameters.

**Table A2:** Physics submodel parameters.

| Parameter Name | Variable Type | Meaning |
| --- | --- | --- |
| substeps | Integer | Number of times the physics algorithm is run. |
| friction | Float | Dampening multiplier for cell velocity, applied each step. |
| cell-radius | Float | Cell radius. |
| cell-repulsion | Float | Repulsion multiplier for cell-cell collisions. |

##### Locomotion

Cell locomotion is modeled by a simplified Correlated Random Walk (CRW) [3, 5] approach. CRWs involve a ‘persistence’ [18] over time in the orientation of successive steps, creating a directional bias in the short term and a uniform distribution in the long term. Our simplified method uses a discrete turning state and stochastic updates to that state to interpolate between linear and curved motion. This method minimally models persistence according to the following equations.

Let each cell have the following properties:

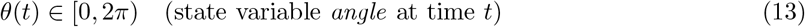

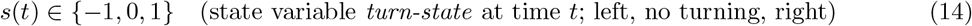

The turn state *s*(*t*) is updated each time step according to the following rule:

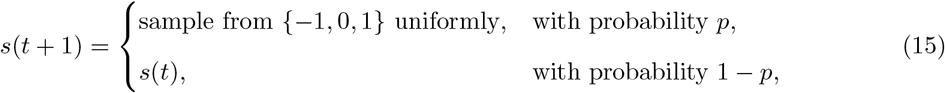

where *p* is submodel parameter *turn-chance*. If the random roll with probability *p* succeeds, the cell updates its turning state to a random choice of left, straight, or right.

Each time step, the cell’s movement angle is incremented by a constant turning speed:

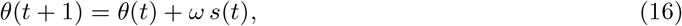

where *ω* is submodel parameter *turn-speed*. The base movement vector of a cell is then given by:

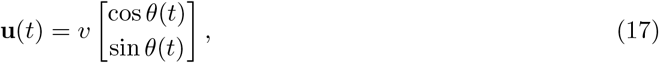

where *v* is the base movement speed. This vector is accumulated in Δ**x**(*t*) in the ECM Deposition submodel to incorporate the effect of ECM on locomotion speed.

In Table A3 we show the locomotion submodel parameters.

**Table A3:** Locomotion submodel parameters.

| Parameter Name | Variable Type | Meaning |
| --- | --- | --- |
| max-speed | Float | Maximum cell speed. |
| turn-speed | Float | Turning speed in radians/step. |
| turn-chance | Float | Chance per step to change state variable *turn-state*. |

##### Cell Cycle

Progression through the cell cycle follows a deterministic approach, for the simplicity of the model. Each cell is assigned a total cell cycle duration T (in simulation steps), which is divided into fixed fractions corresponding with each phase of the cell cycle G1, S, G2, M. This method allows for T to be varied without changing the relative durations of each phase.

Let *τ* ∈ [0, *T*) denote the time elapsed since the start of the current cycle. Cell phase transitions occur when *τ* exceeds the cumulative fraction corresponding to the current phase. During early G1, cells enter G0 if state variable *inhibition-amount* exceeds a configured threshold.

Cell movement speed depends on the current cell cycle phase. Cells in G0 are immobile. Cells in G2 exhibit reduced movement speed in preparation for mitosis, while cells in G1 return to the default movement speed.

When a cell completes M phase, it divides, creating a new cell at an adjacent position with stochastically determined state-variables. Cell cycle time *τ* is reset for both cells.

In Table A4 we show the cell cycle submodel parameters.

**Table A4:** Cell cycle submodel parameters.

| Parameter Name | Variable Type | Meaning |
| --- | --- | --- |
| avg-cycle-duration | Integer | Default cell cycle duration. |
| mitosis-offset | Float | Distance offset each child cell is placed upon mitosis. |
| inhibition-radius | Float | Radius of neighbor detection (to count towards contact inhibition). |
| inhibition-increment | Float | Amount by which state variable *inhibition-amount* is incremented for each neighbor cell. |

##### ECM Deposition

Every time step, the cell is repulsed by the centroid of nearby ECM particles within a radius. The equations below describe the mechanics of this repulsion. The set of nearby ECM particles is defined as:

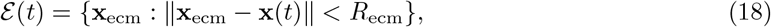

where *R*_*ecm*_ is submodel parameter *ecm-detection-radius*. The centroid of nearby ECM is defined by

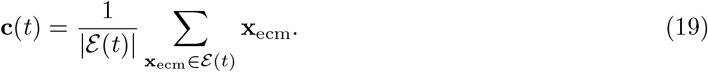

An ECM bias vector is then calculated which directs the cell down the concentration gradient of ECM particles:

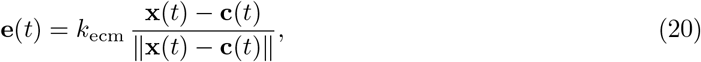

where *k*_*ecm*_ is submodel parameter *ecm-avoidance*. Finally, the ECM speed reduction scalar *w* is calculated by

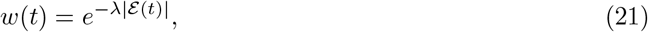

where *λ* is submodel parameter *ecm-speed-scalar*. This scalar slows cell-driven movement when surrounded by high concentrations of ECM. It is incorporated in both the locomotion and ECM deposition position updates, according to the following equations:

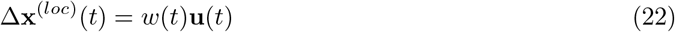

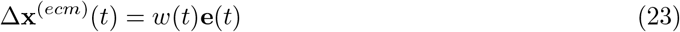

Finally, **x**(*t*) is updated as described in (7) and (8).

In Table A5 we show the ECM deposition submodel parameters.

**Table A5:**
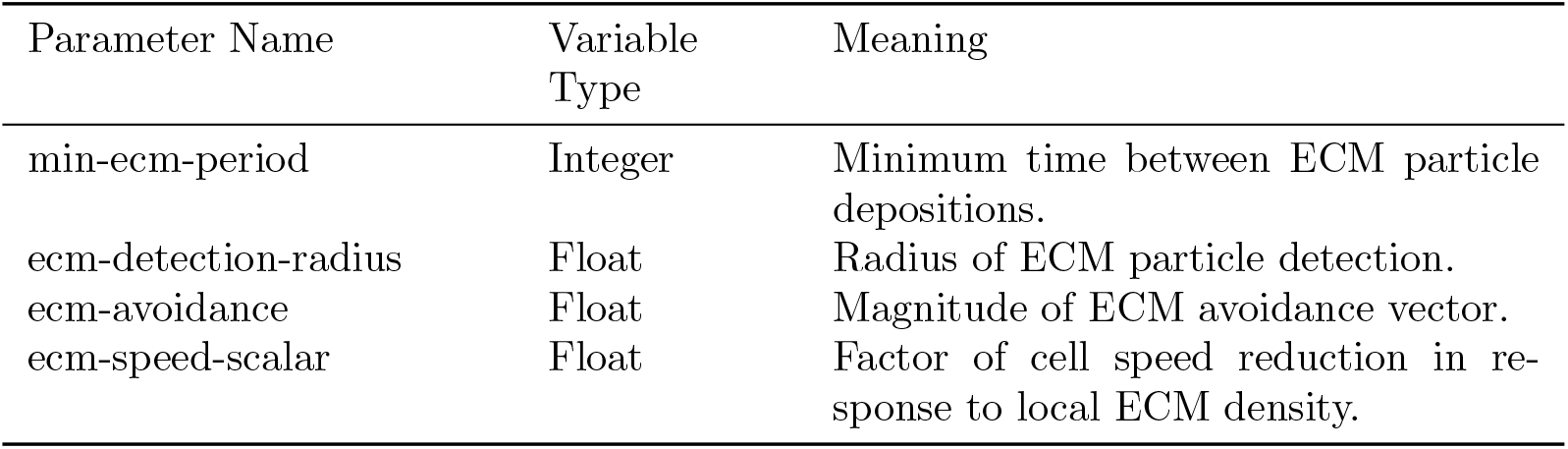
ECM Deposition submodel parameters.

| Parameter Name | Variable Type | Meaning |
| --- | --- | --- |
| min-ecm-period | Integer | Minimum time between ECM particle depositions. |
| ecm-detection-radius | Float | Radius of ECM particle detection. |
| ecm-avoidance | Float | Magnitude of ECM avoidance vector. |
| ecm-speed-scalar | Float | Factor of cell speed reduction in response to local ECM density. |

## References

[1] Julia C. Arciero, Qi Mi, Maria F. Branca, David J. Hackam, and David Swigon. Continuum model of collective cell migration in wound healing and colony expansion. Biophysical Journal, 100(3):535–543, Feb 2011.

[2] Aikaterini Berdiaki, Monica Neagu, Petros Tzanakakis, Ioanna Spyridaki, Serge Pérez, and Dragana Nikitovic. Extracellular matrix components and mechanosensing pathways in health and disease. Biomolecules, 14(9), 2024.

[3] Pierre Bovet and Simon Benhamou. Spatial analysis of animals’ movements using a correlated random walk model. Journal of Theoretical Biology, 131(4):419–433, 1988.

[4] Francesca Campolongo, Jessica Cariboni, and Andrea Saltelli. An effective screening design for sensitivity analysis of large models. Environmental Modelling & Software, 22(10):1509–1518, 2007.

[5] Edward A. Codling, Michael J. Plank, and Simon Benhamou. Random walk models in biology. Journal of the Royal Society, Interface, 5(25):813–834, 2008.

[6] Jillian Cwycyshyn, Cooper Stansbury, Walter Meixner, James B. Hoying, Lindsey A. Muir, and Indika Rajapakse. A programmable platform for probing cell migration and proliferation. APL Bioengineering, 8(4):046106, 2024.

[7] John C. Dallon, Jonathan A. Sherratt, and Philip K. Maini. A mathematical model of the effects of growth factors on dermal wound healing. Mathematical Biosciences, 162(1–2):157–177, 1999.

[8] R. B. Dickinson, S. Guido, and R. T. Tranquillo. Biased cell migration of fibroblasts exhibiting contact guidance in oriented collagen gels. Annals of Biomedical Engineering, 22(4):342–356, July 1994.

[9] F. Graner and J. A. Glazier. Simulation of biological cell sorting using a two-dimensional extended potts model. Physical Review Letters, 69(13):2013–2016, Sep 1992.

[10] Volker Grimm, Uta Berger, Finn Bastiansen, Sigrunn Eliassen, Vincent Ginot, Jarl Giske, John Goss-Custard, Tamara Grand, Simone K. Heinz, Geir Huse, Andreas Huth, Jane U. Jepsen, Christian Jørgensen, Wolf M. Mooij, Birgit Müller, Guy Pe’er, Cyril Piou, Steven F. Railsback, Andrew M. Robbins, Martha M. Robbins, Eva Rossmanith, Nadja Rüger, Espen Strand, Sami Souissi, Richard A. Stillman, Rune Vabø, Ute Visser, and Donald L. DeAngelis. A standard protocol for describing individual-based and agent-based models. Ecological Modelling, 198(1):115–126, 2006.

[11] Volker Grimm, Uta Berger, Donald L. DeAngelis, J. Gary Polhill, Jarl Giske, and Steven F. Railsback. The odd protocol: A review and first update. Ecological Modelling, 221(23):2760–2768, 2010.

[12] Volker Grimm, Steven F. Railsback, Christian E. Vincenot, Uta Berger, Cara Gallagher, Donald L. DeAngelis, Bruce Edmonds, Jiaqi Ge, Jarl Giske, Jürgen Groeneveld, Alice S.A. Johnston, Alexander Milles, Jacob Nabe-Nielsen, J. Gareth Polhill, Viktoriia Radchuk, Marie-Sophie Rohwäder, Richard A. Stillman, Jan C. Thiele, and Daniel Ayllón. The odd protocol for describing agent-based and other simulation models: A second update to improve clarity, replication, and structural realism. Journal of Artificial Societies and Social Simulation, 23(2):7, 2020.

[13] Max D. Morris. Factorial sampling plans for preliminary computational experiments. Technometrics, 33(2):161–174, 1991.

[14] Matthias Müller, Bruno Heidelberger, Marcus Hennix, and John Ratcliff. Position based dynamics. J. Vis. Comun. Image Represent., 18(2):109–118, April 2007.

[15] Fumihiko Nakamura. The role of mechanotransduction in contact inhibition of locomotion and proliferation. International Journal of Molecular Sciences, 25(4):2135, Feb 2024.

[16] Lars Olsen, Jonathan A. Sherratt, and Philip K. Maini. A mechanochemical model for adult dermal wound contraction: On the permanence of the contracted tissue displacement profile. Journal of Theoretical Biology, 177(2):113–128, 1995.

[17] Anna-Maria Pabst, Ulrich Hiden, Andreas König, Julia König, Christine König, et al. Human in vitro skin models for wound healing and wound healing disorders. Biomedicines, 11(4):1056, 2023.

[18] Clifford S. Patlak. Random walk with persistence and external bias. The Bulletin of Mathematical Biophysics, 15(3):311–338, 1953.

[19] Reza Riahi, Yongliang Yang, Donna D. Zhang, and Pak Kin Wong. Advances in wound-healing assays for probing collective cell migration. Journal of Laboratory Automation, 17(1):59–65, 2012.

[20] Robert T. Tranquillo and J. D. Murray. Continuum model of fibroblast-driven wound contraction: Inflammation-mediation. Journal of Theoretical Biology, 158(2):135–172, 1992.

[21] D. C. Walker, G. Hill, S. M. Wood, R. H. Smallwood, and J. Southgate. Agent-based computational modeling of wounded epithelial cell monolayers. IEEE Transactions on Nanobioscience, 3(3):153–163, 2004.

